# Assessing the Effectiveness of 3D-Printed Ceramic Structures for Coral Restoration: Growth, Survivorship, and Biodiversity Using Visual Surveys and eDNA

**DOI:** 10.1101/2025.06.23.660499

**Authors:** Vriko Yu, Alison Corley, Horace Lau, Philip D. Thompson, Zhongyue Wilson Wan, Jane C.Y. Wong, Shelby McIlroy, David M. Baker

## Abstract

Coral reef degradation has spurred the development of artificial structures to mitigate losses in coral cover. These structures serve as substrates for coral transplantation, with the expectation that growing corals will attract reef-associated taxa — while the substrate’s ability to directly support biodiversity is often neglected. We evaluated a novel 3D-printed modular tile made of porous terra cotta, designed with complex surface structures to enhance micro- and cryptic biodiversity, through a restoration project in Hong Kong. Over four years, we monitored 378 outplanted coral fragments using diver assessments and photography, while biodiversity changes were assessed through visual surveys and eDNA metabarcoding. Coral survivorship was high, with 88% of transplants surviving by the study’s end. The restoration site exhibited greater fish and macroinvertebrate abundance compared to a nearby unrestored area. eDNA analyses revealed 23.5% higher eukaryote ASV richness at the restoration site than the unrestored site and a 13.3% increase relative to a natural reference coral community. This study highlights the tiles’ dual functionality: (1) supporting coral growth and (2) enhancing cryptic biodiversity, an aspect often neglected in traditional reef restoration efforts. Our findings underscore the potential of 3D-printed ceramic structures to improve both coral restoration outcomes and broader reef ecosystem recovery.

## 1. Introduction

Coral reefs are declining at an alarming rate. Driven by the combined impacts of climate change, overfishing, eutrophication, and coastal development, coral cover worldwide has already declined by over 50% since the late 20th century, with some regions — such as the Caribbean — experiencing losses of more than 80% [1–4]. As mass bleaching events and coral mortality become more frequent and intense, over 90% of the world’s reefs are projected to face severe degradation by the middle of the century [5,6]. This collapse threatens not only marine biodiversity — with up to a quarter of all marine species depending on coral reefs [7,8]— but also the livelihoods of roughly 500 million people and ecosystem services valued at up to USD 350,000 per hectare annually [9,10].

In response, Artificial Reefs (ARs) have emerged as one of the most widely used interventions to stabilize degraded reef structures, promote coral settlement, and support broader ecological recovery [11,12]. ARs range from simple rubble piles and concrete blocks to complex, biomimetic structures designed to increase surface area, reduce hydrodynamic stress, and provide microhabitats for diverse marine organisms [13]. When combined with coral outplanting, ARs can enhance survivorship and potentially accelerate the recovery of benthic communities. However, many ARs remain reliant on non-biocompatible materials, provide minimal fine-scale habitat complexity, are unsuited for sediment-laden or high-energy environments, and are limited in their scalability for covering large areas. In addition, few artificial structures produced for coral reef restoration explicitly consider cryptic biodiversity or long-term ecological function in their design, with most focusing solely on mitigating losses in coral cover [12].

To address these limitations, we developed a ceramic 3D-printed reef tile, engineered from terracotta clay and designed to integrate both coral restoration and biodiversity enhancement through a scalable, modular design. The tiles feature a hexagonal configuration and fine-scale topography intended to promote coral attachment, reduce sediment smothering, and replicate natural reef crevices, thus facilitating larval recruitment and benthic colonization [14,15]. Unlike conventional concrete ARs, the reef tiles are deployable directly onto soft benthic substrates without complex substrate engineering, offering a low-impact solution for challenging or urbanized environments. The material properties of terracotta — being porous, erosion-resistant, and biocompatible — further support microbial colonization and ecological succession, enhancing both coral health and overall habitat function [16,17]. Yet, the effectiveness of these design choices requires empirical validation. Beyond coral fragment survivorship, the potential of the artificial structures and the outplanted coral community to facilitate the reassembly of reef-associated taxa — which are critical for restoring ecological function — must also be assessed under dynamic, real-world conditions.

While coral survivorship remains a central restoration target, ecosystem resilience depends equally on the reassembly of diverse coral-associated communities that sustain trophic interactions and contribute to reef function [11,18–20]. Owing to their relative ease of identification, sensitivity to environmental disturbance, and role in mitigating macroalgal overgrowth, macroinvertebrates and fish are often the only targets for biodiversity monitoring in coral restoration projects [21]. Beyond these two taxonomic groups however, are a range of organisms that provide equally critical functions on coral reefs: Crustose coralline algae provide larval settlement cues, sponges consolidate loose sediments and rubble, smaller cryptic invertebrates also feed on algae and act as a food source for reef fish — while all of these examples, in combination with the microbial communities of bacteria, algae, and fungi, facilitate nutrient cycling [22].

To capture a fuller picture of the effect of the artificial reef tiles on community diversity, we combined standard SCUBA-based visual surveys of fish and macroinvertebrates with environmental DNA (eDNA) metabarcoding, a non-invasive tool that enables detection of both conspicuous and cryptic taxa from environmental samples [23,24]. eDNA is a particularly useful tool for monitoring structurally complex systems like coral reefs, where its ability to detect a wide range of organisms complements traditional visual surveys and bridges critical gaps in current restoration monitoring frameworks [25]. In this study, we evaluated the ecological performance of 3D-printed ceramic reef tiles deployed in a subtropical, urbanized marine environment to determine whether this engineered substrate can support scalable coral restoration while promoting broader biodiversity uplift with the goal of providing a replicable model for effective, ecosystem-based reef rehabilitation.

## 2. Materials and Methods

### 2.1. Study Sites

Lying just south of the Tropic of Cancer, Hong Kong supports a surprising level of coral diversity, with more than 90 species of reef-building corals recorded across its semi-enclosed, estuarine-influenced waters [26–28]. This diversity is particularly noteworthy, given the region’s marked seasonal variability and persistent anthropogenic stressors, such as coastal development, salinity fluctuations, and sediment-laden runoff [29]. Coral cover in Hong Kong declined substantially over the past century, driven by acute stressors such as thermal anomalies and extreme weather events, alongside chronic local pressures like nutrient enrichment and elevated suspended solids [30,31]. Since the 1990s, however, water quality in Hong Kong has improved due to stricter controls on wastewater treatment [32]. Hong Kong thus presents a suitable environment to explore the viability of re-introducing extirpated corals following a reduction in the extent of local environmental stressors [26].

This study was conducted within Hoi Ha Wan Marine Park (HHWMP; 22°28′13.68″N, 114°20′08.25″E), located in the northeastern waters of the Sai Kung Peninsula, Hong Kong SAR (Figure 1). Established in 1996, HHWMP spans approximately 260 hectares and is known for its diverse subtropical coral communities, supporting more than 60 species of hard corals (AFCD Report, 2004). The park experiences pronounced seasonal variation, with sea surface temperatures ranging from 15 °C in winter to 28 °C in summer, and salinity modulated by monsoonal rainfall and freshwater inflow. These dynamic conditions, combined with proximity to urbanized coastal zones, positions HHWMP as a representative site for testing coral restoration strategies in marginal, subtropical reef environments.

**Figure 1.**
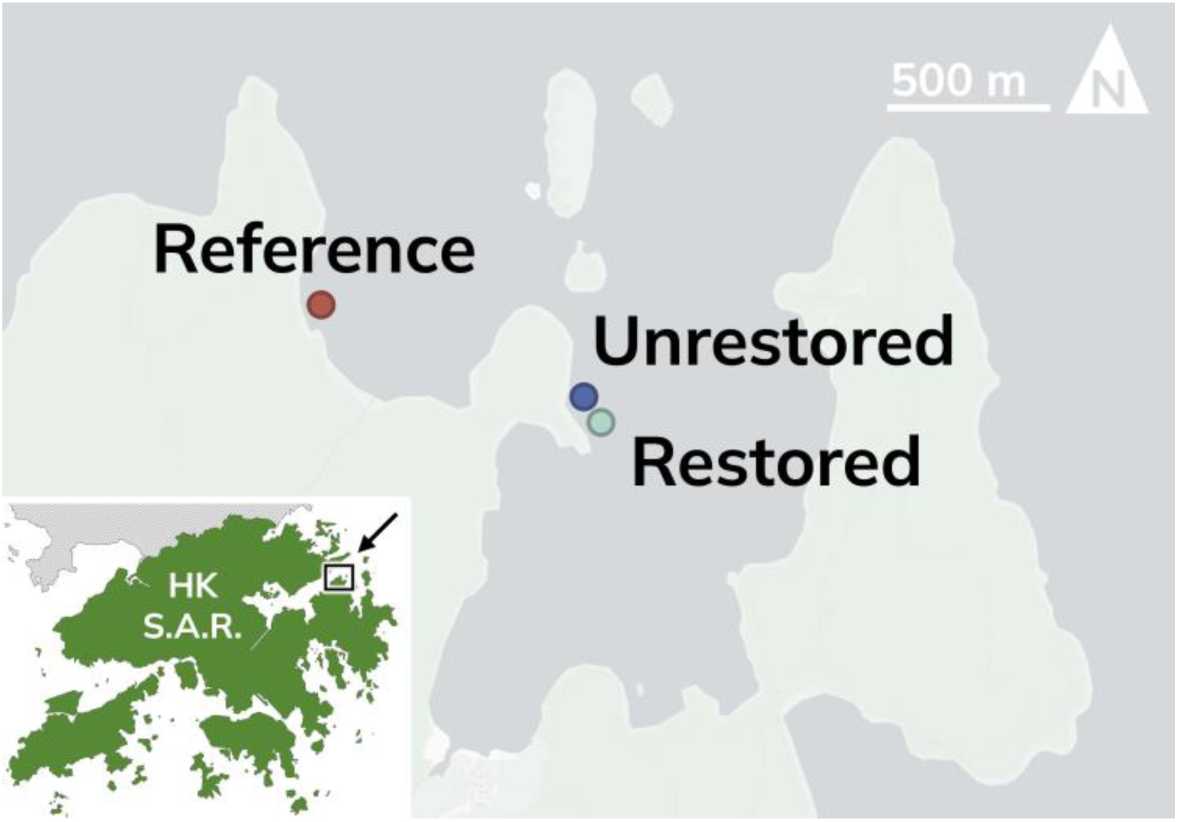
Locations of study sites in Hoi Ha Wan Marine Park, Hong Kong. Map showing the positions of the reference (Gruff Head), restored (Coral Beach), and unrestored (Coral Beach) sites surveyed in this study. Basemap data from © OpenStreetMap contributors and © CARTO, accessed via the QuickMapServices plugin in QGIS.

Three ecologically distinct sites within HHWMP were selected for comparative analysis (Figure 1). The restored site, off Coral Beach, served as the primary deployment area for the reef tiles. Prior to the installation of the artificial structures, this site was a sandy seabed with no coral cover or hard substrate. Approximately 50 m north of the restored area, parallel to the shoreline, the unrestored site comprised a section of sandy seabed with no artificial substrate, representing a degraded benthic baseline with no intervention. The reference site, situated at Gruff Head approximately 500 m west of the restored site, featured a natural coral assemblage and minimal anthropogenic disturbance. This site was selected to represent a realistic restoration target for community diversity and function. All three sites were located at comparable depths (6-8 m) and experienced similar hydrodynamic and environmental conditions.

### 2.2. Reef Tile Design

The 3D-printed ceramic reef tiles used in this study were custom-engineered ceramic substrates designed to promote coral recruitment, enhance structural complexity, and mitigate sediment accumulation in a subtropical marine environment. Each tile was hexagonal in shape, with a maximum width of 650 mm, and comprised two integrated functional layers: a structural base layer (“grid layer”) and a biomimetic surface layer (“coral layer”) (Figure 2). The grid layer featured a diagrid framework with integrated lateral bracing to maximize mechanical stability while minimizing material usage and fabrication defects. The geometry was optimized to reduce cracking during drying and thermal stress during firing, with a print path totaling approximately 170 m in length. This design facilitated uniform moisture evaporation and enhanced compressive strength under load. To improve hydrodynamic performance and minimize sediment deposition, a three-legged footing system was printed on the underside of each tile after drying (Figure 2). These legs elevated the tile slightly above the seabed to promote water flow and create a sheltered interface for benthic recruitment.

**Figure 2.**
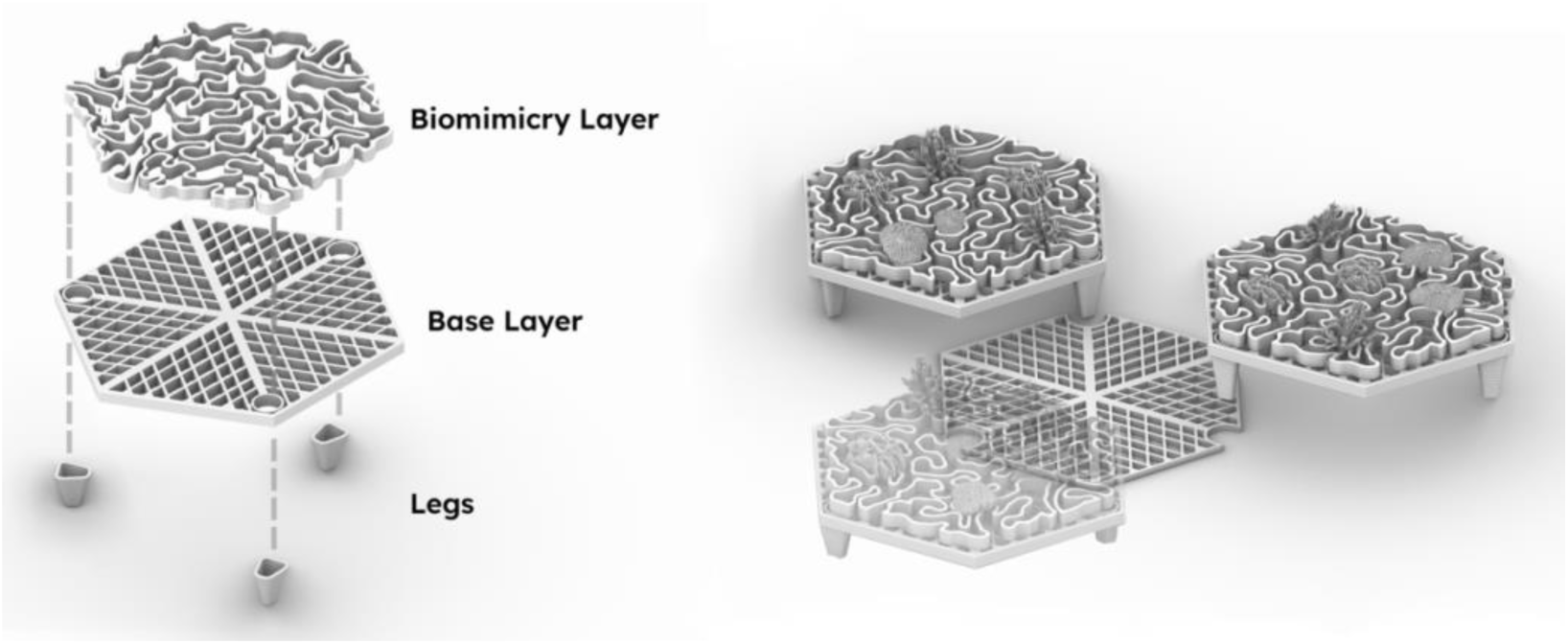
Design and structural components of the custom-engineered 3D-printed reef tiles used for coral recruitment and restoration. (Left) The upper “biomimicry layer” of the tiles features a biomimetic surface designed to mimic reef complexity, providing habitat for cryptic invertebrates. The grid layer has a diagrid framework and lateral bracing to enhance mechanical strength, reduce fabrication defects, and support uniform drying and firing. The integrated three-legged footing system is printed and attached post-processing to elevate the tile above the seabed, improving hydrodynamic flow and minimizing sediment accumulation. (Right) One assembled unit showing three tiles, onto which coral fragments were transplanted, anchored on one Base Layer plate. 24 of these assembled units were deployed in 5 m intervals along a 3x8 unit grid at the restoration Site [33].

Fabrication of the 3D-printed reef tiles was performed using a Direct Ink Writing (DIW) method with an ABB 6700 robotic arm and a Deltabots linear ram extruder (6 mm nozzle diameter). The paste material consisted of red terracotta clay (P1331, Potterycrafts Ltd, UK) amended with <1% fine crystalline silica. Tiles were printed with a layer height of 2.7 mm at extrusion speeds ranging from 10.5 to 17 mm/s depending on local path geometry. After air-drying and structural inspection, the tiles were fired in a gas kiln at 1125 °C, yielding a mean shrinkage of approximately 11%.

### 2.3. Restoration Site Design

The coral restoration experiment was conducted within a 10 m × 35 m plot at the restored site (Coral Beach) in Hoi Ha Wan Marine Park (Figure 3). Within this plot, a total of 24 modular reef units — comprising 72 reef tiles — were deployed in a grid layout with 5 m spacing between units. Each unit consisted of three interlocked tiles arranged in a triangular configuration (Figure 2). This modular design enhanced structural heterogeneity while facilitating standardized monitoring across units.

**Figure 3.**
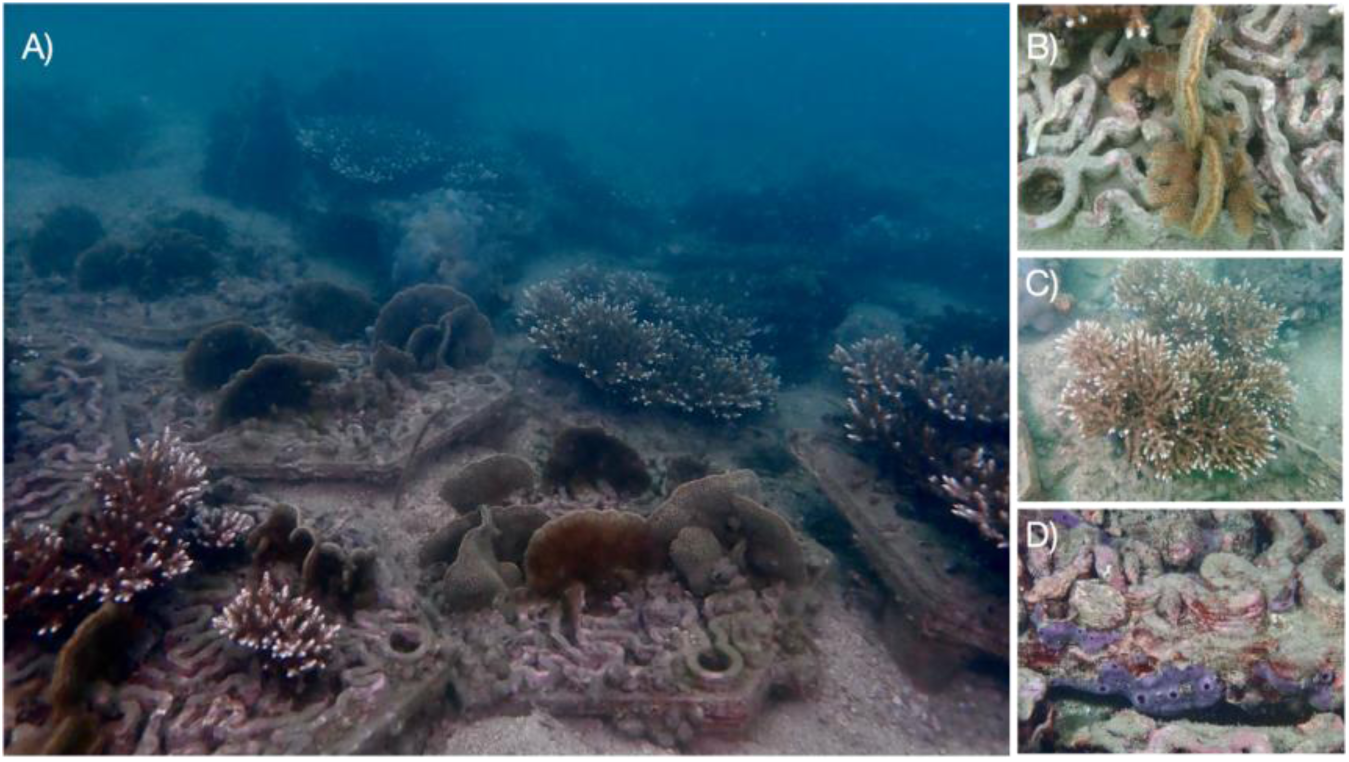
Composite image of the restoration project, 4 years after deployment. (A) A subset of the ceramic reef tiles deployed at the restoration site in Hoi Ha Wan Marine Park, outplanted with multiple coral species representing branching, plating, and massive colony morphologies. (B) Self-attachment of a transplanted fragment via new growth onto the biomimicry layer. (C) A tile seeded with *Acropora* fragments, which have grown to obscure the existence of the reef tile below. (D) Encrusting taxa that have grown onto the terra cotta surface of the reef tile, including marine sponges and coralline algae.

Each reef tile was seeded with six evenly spaced coral fragments, representing three distinct morpho-functional groups: *Acropora* (branching), *Pavona* (plating), and *Platygyra* (massive). *Pavona* and *Platygyra* fragments were collected as Corals of Opportunity within HHWMP, while *Acropora* fragments were sourced from healthy donor colonies at Bluff Island (∼8 km distance from the restoration site), due to their local extirpation within the marine park. A total of 378 fragments were transplanted to the restoration site, consisting of 126 fragments per genus. Coral fragments were secured using marine-grade epoxy (Z-Spar A-788).

### 2.4. Monitoring Coral Survivorship and Growth

Coral performance was assessed quarterly for four years following transplantation, beginning three months after outplanting. SCUBA divers visually examined each coral fragment for signs of bleaching or discoloration at each survey interval. Five conditions were assigned to the coral fragments: healthy, partial mortality, detached, and dead. Corals were considered alive if they were healthy or showed signs of partial mortality; those classified as dead or detached were not counted as having survived. Instances of breakage were also recorded as they occurred. During each monitoring event, high-resolution photographs of each tile and its attached coral fragments were captured using an underwater digital camera (Olympus TG5). Each image included a scale bar placed on the tile, which served as a calibration reference for accurate spatial measurements. Growth measurements were extracted from these photographs using ImageJ software (version 1.52n), recording the maximum linear extension of each coral fragment at each survey date.

Extension rates (cm month^-1^ linear growth) for each coral individual were calculated based on the change in size from the initial measurement to the monitoring survey:

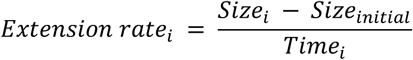

Where *Size*_i_ represents the size at the monitoring survey, *Size_initial_* the initial size measurement, and *Time_i_* the time interval (in months) since the initial measurement. Broken fragments were excluded from growth rate calculations (for all survey dates following the first notice of breakage).

Differences in extension rates and breakage between genera were analyzed using generalized linear mixed models (GLMMs). For extension rates, the model was fitted with the *glmmTMB* package [34], included fixed effects for genus, with a random factor specified for the position of coral fragments nested within transplantation tiles, which were themselves nested within units to capture variability at multiple hierarchical levels. To address temporal autocorrelation among repeated measures on the same coral fragment, a first-order autoregressive (AR1) variance structure was incorporated. Model selection was performed using Akaike’s Information Criterion (AIC) to identify the best-fitting GLMMs (Table A1). Due to the binomial nature of the breakage data, models were fitted using the *lme4* package [35]. When significant differences between genera were detected, pairwise comparisons of estimated marginal means were performed using the emmeans package [36], applying Kenward-Roger degrees-of-freedom method and Tukey-adjusted *p*-values.

### 2.5. Visual Fish and Invertebrate Surveys

To assess reef community’s response to restoration, visual belt transect surveys were conducted concurrently with coral monitoring at all three study sites. At each site, three 35-meter transects were laid out parallel to the shoreline, spaced 5 m apart. Two divers surveyed each transect simultaneously (one on each side), covering a total area of 2 × 35 m per transect. Target species (Table 1) were selected from the list of Reef Check bioindicator species for Hong Kong, which identifies common fish and invertebrate that are used to represent key functional groups and responses to different anthropogenic stressors acting on subtropical reef ecosystems [37]. The monitored species included herbivores and consumers belonging to a range of trophic levels.

**Table 1.**
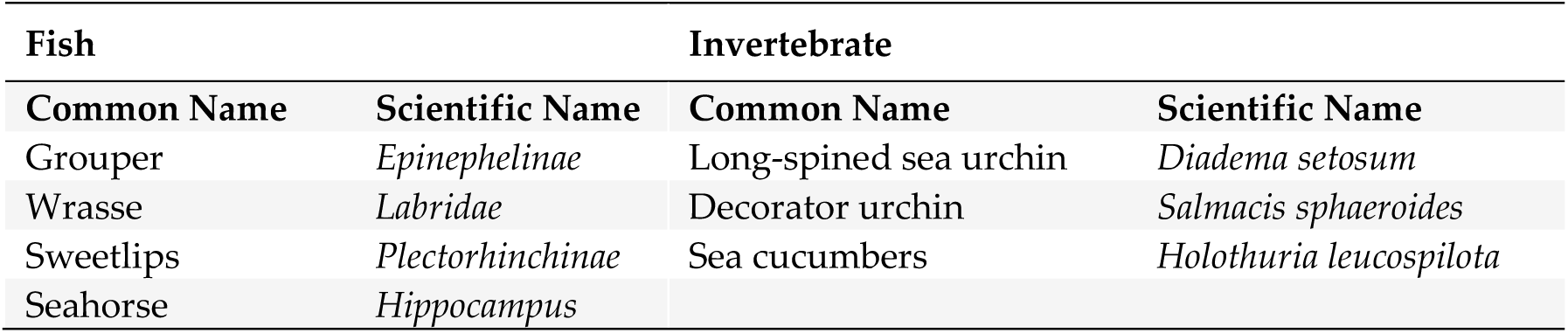
Bioindicator fish and invertebrate species monitored during diver surveys.

| Fish |  | Invertebrate |  |
| --- | --- | --- | --- |
| Common Name | Scientific Name | Common Name | Scientific Name |
| Grouper | *Epinephelinae* | Long-spined sea urchin | *Diadema setosum* |
| Wrasse | *Labridae* | Decorator urchin | *Salmacis sphaeroides* |
| Sweetlips | *Plectorhinchinae* | Sea cucumbers | *Holothuria leucospilota* |
| Seahorse | *Hippocampus* |  |  |

### 2.6 Environmental DNA Metabarcoding

#### 2.6.1 DNA Sampling, Extraction, Amplification, and Sequencing

Two years after the artificial tile deployment, sedimentary environmental DNA (eDNA) was sampled from all three study sites. Five replicates were collected at 5-6 m depth from each site, with 10 m distance between each replicate. Each replicate was collected in a sterile 50 mL falcon tube, collecting sediment from only the top 1 cm of the seafloor. Samples were transported back to the laboratory on ice and stored at −20 °C within three hours of collection. Each sample was homogenized using a sterile spatula and plastic tray, and a 500 mg aliquot of bulk sediment weighed for extraction. DNA was extracted using the DNeasy PowerMax Soil Kit following the manufacturer’s protocol and subsequently purified using the DNeasy PowerClean Cleanup Kit (Qiagen). The concentration of the resulting DNA extracts was quantified using the Qubit dsDNA High Sensitivity Assay (Thermo Fisher Scientific).

Extracted DNA was amplified using the mlCOIintF and jgHCO2198 primer set, which amplifies a ∼313bp fragment of the cytochrome c oxidase subunit I (COI) gene to target metazoan diversity. Polymerase chain reactions (PCR) were prepared to a 20 uL total reaction volume. The reaction mixture included 2 µL of 10X Advantage 2 PCR Buffer (Takara Bio), 0.4 µL of 50X Advantage 2 Polymerase Mix, 1.4 µL of a 50X dNTP Mix (10 mM each), 1 µL of each primer (10 pmol/µL), 0.5 μL Bovine Serum Albumin (20 mg/mL), and 10 to 30 ng of template DNA. The following thermocycling parameters were used: an initial 1 minute at 95 °C, followed by 16 cycles of denaturing at 95 °C for 10 seconds, annealing at 62 °C for 30 seconds, and extension at 72 °C for 1 minute, ending with 10 minutes of final extension at 72 °C. To increase the probability of amplification for low-copy DNA sequences, DNA extracted from each sample was amplified in triplicate. At minimum, one PCR control was performed per seven sample reactions using autoclaved MQ as the template. PCR products were visualised via electrophoresis in a 2% agarose gel. Triplicate amplification products for each sample were then pooled and subsequently purified using the QIAquick Gel Extraction Kit (Qiagen). A unique 6 bp indexing primer was attached to each pooled sample.

Library preparation and sequencing was completed by Genewhiz (Suzhou, China). All 15 samples submitted for sequencing passed initial quality control, with DNA concentrations ranging between 0.2 and 23.4 ng/μL. Sequencing was performed on an Illumina MiSeq system. A total of 24.6 million 251 bp paired-end reads were generated, with 1.4 to 1.8 million reads per sample. A binned quality score of Q30 (indicating base-call accuracy of 99.9%) was given to 90% of base calls.

#### 2.6.2. Bioinformatics

##### 2.6.2.1 Sequence Analysis and Quality Control

Raw sequencing reads were processed in *QIIME2* (v2022.2) [38] using the *DADA2* plugin (v1.22) [39]. Reads were demultiplexed in *QIIME2*. The *denoise-paired* function of *DADA2* was used to trim primers, denoise and join paired reads, dereplicate duplicate sequences, and remove chimeric sequences. Forward and reverse reads were both trimmed by 26 bp at the 5’ end and truncated at the 3’ end at 250 bp. Chimeras were removed based on the consensus method. A total of 117,673 amplicon sequence variants (ASVs), represented by 18,704,812 reads, were inferred. Taxonomy was then assigned using the Ribosomal Database Program (*RDP*) classifier v2.13 [40], with a pre-trained database of *COI* reference sequences (v4) [41]. Taxonomic nomenclature was manually edited as needed to adhere to the terminology established by the World Register of Marine Species (*WoRMS*; www.marinespecies.org).

Following taxonomic assignment, data were imported into *R* (v4.2.1) [42]; R Core Team, 2018) for further filtering and analysis using the *phyloseq* package (v1.38) [43]. Alpha rarefaction curves were plotted using the *ggrare()* function of the *ranacapa* package (v0.1) [44]. One sample from the unrestored site had anomalously low ASV richness (974 ASVs) relative to other samples from the same site (4,813 to 16,559 ASVs per sample; Figure 2), with the overwhelming majority of reads from this sample (>95.6%) assigned to only two species. This sample did not meet quality standards and was removed from the dataset. Singletons (1,050 ASVs) were then removed, and sequences assigned to Bacteria (23,568 ASVs), Archaea (3 ASVs), or terrestrial taxa (6,431 ASVs) were discarded.

Stringent prevalence and abundance filters were applied to eliminate spurious ASVs produced by background sequencing errors during apparent over-sequencing (Figure A1). Only ASVs that appeared in at least three samples from each site were retained. To mitigate bias in prevalence filtering introduced by unequal replication between sites, a mock sample was generated by randomly sampling from a combined pool of ASVs from all other unrestored site samples (seed value = 852). The prevalence filter removed 94.8% of the remaining ASVs (81,583 ASVs), which accounted for 41.8% of the remaining sequencing reads. ASVs were then filtered based on abundance, removing those with a cumulative read abundance of fewer than 50 total counts across all samples (493 ASVs). Samples were then rarefied to an even sequencing depth (96,443 reads per sample, 90% of the depth of the sample with the lowest number of reads; Figure A1). No ASVs were removed through rarefaction.

##### 2.6.2.2. Community Diversity Analysis

All analyses were performed in *R* using *phyloseq* (v1.38) [43]. Figures were created using *ggplot2* (v3.5) [45]. A tandem approach was adopted to examine taxonomic composition and differences in community structure between sites, using both occurrence and relative read abundance data. Two transformed ASV tables were utilized: one in which read counts were converted to a binary presence-absence matrix, and another in which read counts were retained and normalized through Hellinger transformation. For metrics and visualizations based on the site-level eDNA community, ASV counts for each replicate were merged by site prior to transformation.

Observed ASV richness was used to measure alpha diversity, comparing both the number of unique ASVs detected across all samples for each site and the ASV richness of each replicate by site. To test whether there was a significant difference in sample ASV richness between sites, a one-way analysis of variance (ANOVA) was conducted with the *vegan* package (v2.5) [46]. Homogeneity of variance and normality were verified by Levene’s and Shapiro’s tests using the *rstatix* package (v0.7) [47]. The ASV richness of the 10 most diverse phyla by site was visualized as a stacked bar plot, with the number of ASVs among the remaining, less-represented phyla pooled into one group. A chi-squared contingency table test was used to determine whether ASV richness for individual phyla varied significantly by site, using the *chisq.test()* function of the *stats* package. To minimize statistical bias introduced by uneven ASV richness across different phyla, only phyla represented by at least 50 ASVs were included in the chi-squared test, and two versions of the test were conducted: one including Bacillariophyta, which had a significantly higher number of ASVs compared to all other taxa, and another excluding Bacillariophyta.

Beta diversity was assessed using two dissimilarity matrices: one based on the Jaccard index, calculated from occurrence data only, and another based on Bray-Curtis dissimilarity, calculated from Hellinger-transformed read abundances. Principal coordinate ordinations (PCoA) based on each of these matrices were produced to visualize differences in community composition between samples. Differences between sites were then tested using permutational analyses of variance (PERMANOVA) on both matrices with the *adonis2* function of *vegan*. This was followed by post-hoc pairwise comparisons using the *pairwiseAdonis* wrapper for *vegan [48]*. Taxonomic composition plots were created from the Hellinger-transformed read count matrix. Three plots were generated: (1) total Hellinger-transformed read abundance by sample, to better understand variation in sample composition among replicates from the same site; (2) compositional abundance by site, where sample data were merged, Hellinger-transformed, and then converted to relative abundance data to approximate a consensus composition for each site; and (3) another version of the same site-level compositional abundance, but with Bacillariophyta sequences removed prior to data transformation, to better examine differences in the relative abundance of less-abundant phyla.

## 3. Results

### 3.1. Coral Performance

#### 3.1.1. Coral Transplant Survival

Survivorship of transplanted coral fragments remained high throughout the four-year monitoring period, with 88% of individuals (n = 378) classified as alive (86% healthy, 2% partial mortality) and securely attached at the study’s conclusion (Figure 4; Table A2). Full mortality occurred in just 2% of fragments, while 10% were lost due to detachment from the reef tile (Table A2). Mortality and detachment rates remained low and stable over time, indicating sustained substrate stability and fragment viability at the restoration site. No bleaching events were recorded during the monitoring surveys.

**Figure 4.**
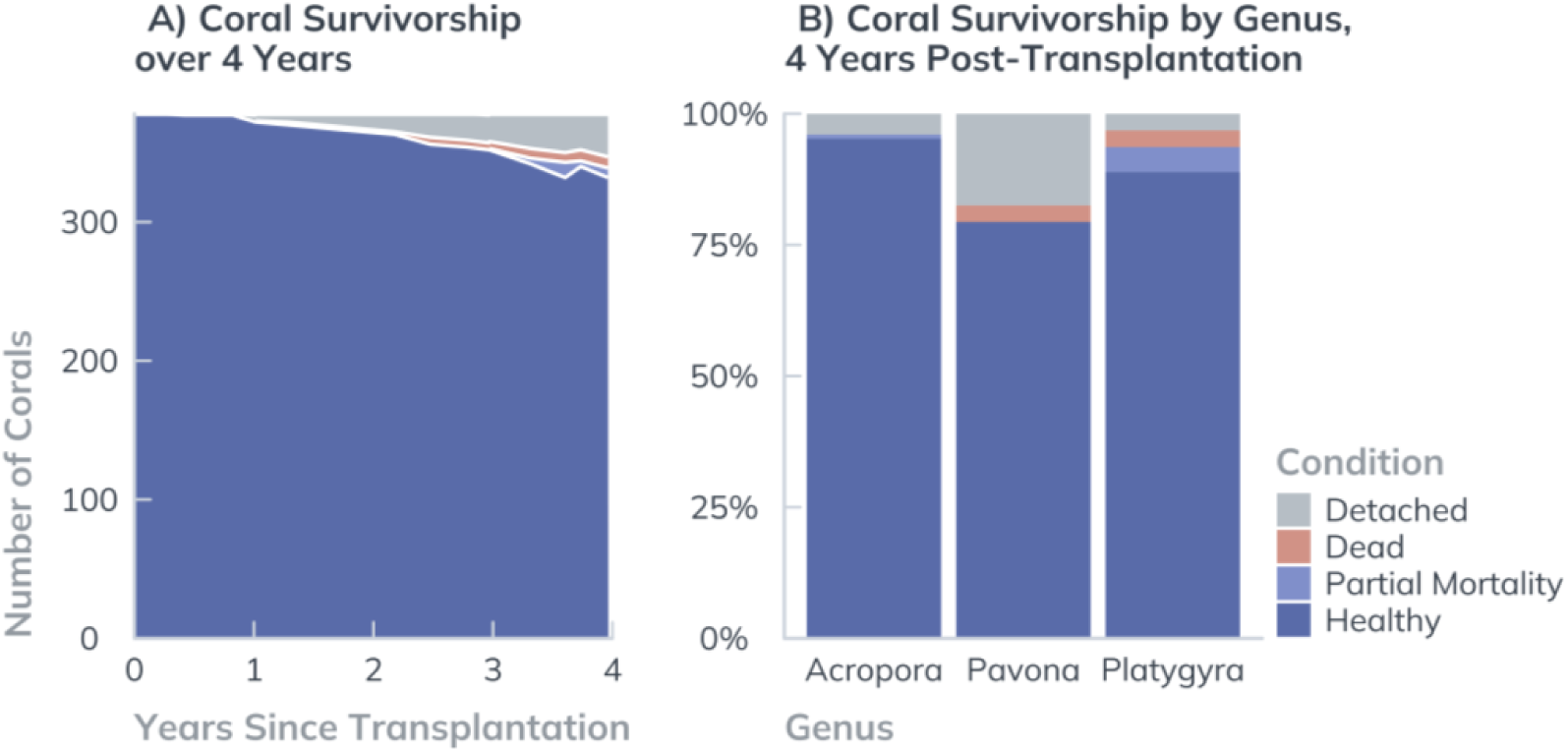
Summary of coral survivorship and condition. (A) The overall survivorship of corals, and (B) genus-specific survivorship of *Acropora, Pavona*, and *Platygyra* over a 4-year period following transplantation. Different conditions were recorded as Healthy (blue), Partial Mortality (light blue), Dead (red), and Detached (grey).

Genus-level analysis revealed variation in resilience among the three coral growth morphologies (Figure 4). *Acropora* exhibited the highest survivorship, with 94% of fragments remaining healthy and only 1% showing partial tissue loss. Of the 6% of *Acropora* fragments that were lost, all were due to detachment. *Platygyra* fragments performed less well, with 89% of fragments classified as healthy. Furthermore, 5% exhibited partial mortality, 3% were dead, and an additional 3% had detached after four years. *Pavona* had the lowest survivorship rate, with only 76% classified as healthy, 3% dead, and 21% detached by the end of the monitoring period.

#### 3.1.2. Extension Rates and Breakage

Over the four-year monitoring period, all three coral genera exhibited positive linear extension relative to their initial transplantation size, though significant differences in growth trajectories were observed among genera representing distinct growth forms (Figure 5). *Acropora* fragments showed the most pronounced increase in linear extension, with a mean growth of 254% relative to baseline measurements, corresponding to an extension rate of 0.35 ± 0.16 cm month^-1^ (*n* = 1495). *Pavona* displayed intermediate growth, achieving a mean increase of 179%, with an extension rate of 0.20 ± 0.11 cm month^-1^ (*n* = 1295). In contrast, *Platygyra*, characterized by its massive growth form and inherently slower growth dynamics, with a mean increase of 118% and an extension rate of 0.14 ±

**Figure 5.**
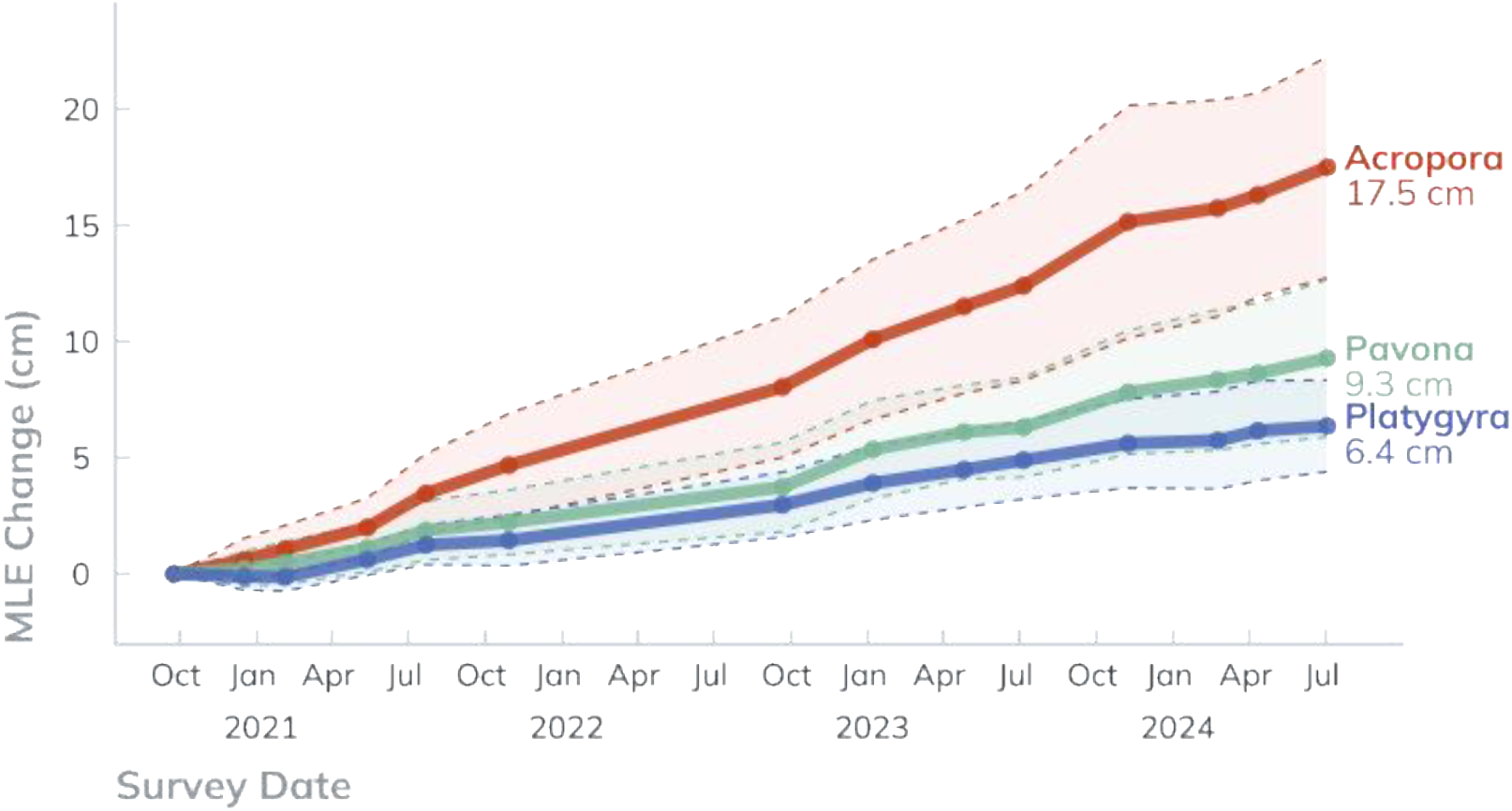
Coral Growth from transplantation size. Change in maximum linear extension (MLE, cm) of the three genera of corals, *Acropora* (red), *Pavona* (green), and *Platygyra* (blue) over a 4-year period following transplantation. The solid line represents the mean values; the dotted lines indicate standard deviation.

0.06 cm month^-1^ (*n* = 1335). Tukey-adjusted contrasts of growth rates (Table 2) showed that *Acropora* grew significantly faster than *Pavona* (z = 10.482, *p* < 0.001) and *Platygyra* (*z* = 19.071, *p* < 0.001). *Pavona* also grew faster than *Platygyra* (*z* = 8.030, *p* < 0.001). The incidence of broken fragments was also examined (Table 2), with the highest in *Acropora*, significantly more than *Platygyra* (z = 7.0899, *p* < 0.001) and *Pavona* (z = 7.472, *p* < 0.001).

**Table 2.**
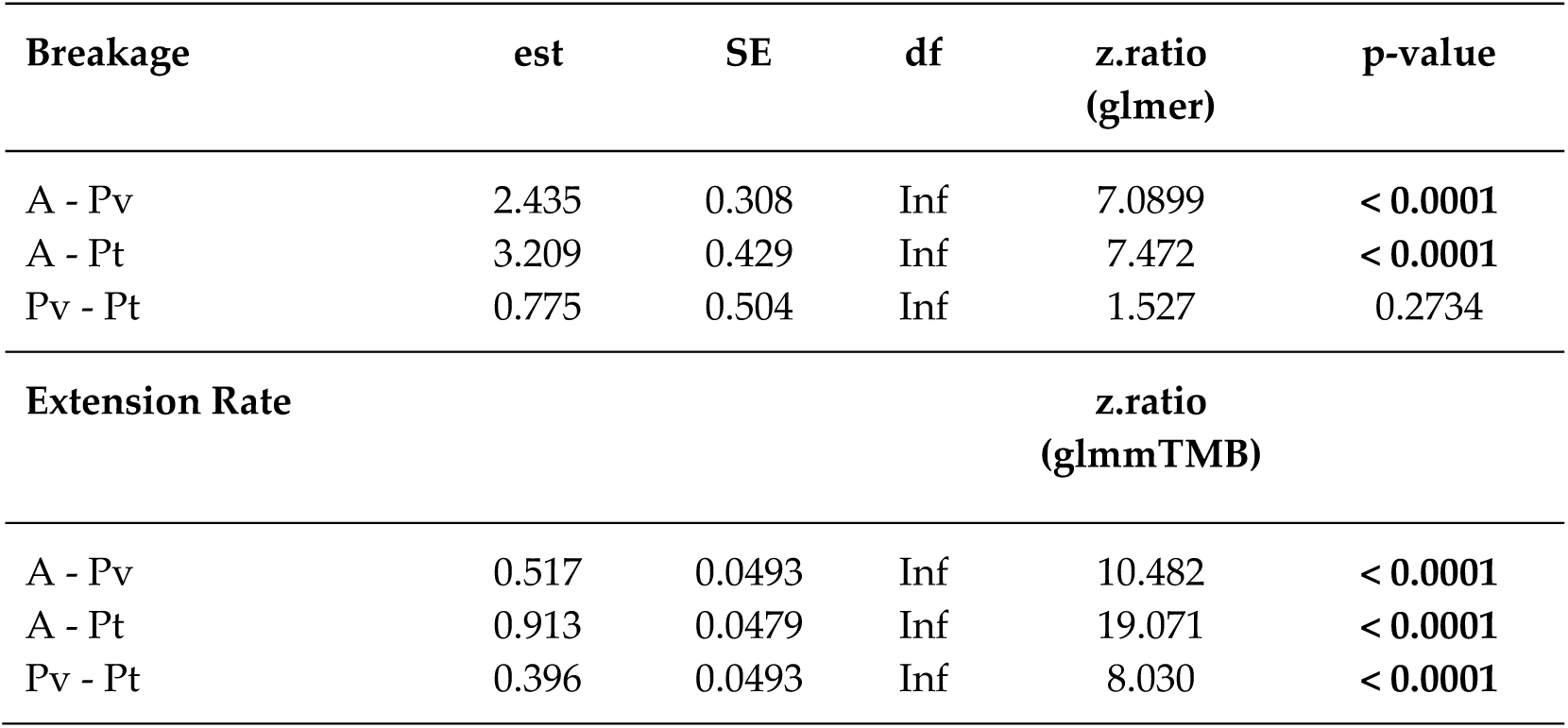
Results of the GLMM model and pairwise comparisons of estimated marginal means (EMMs) examining the effect of coral genera on extension rate and breakage. *p*-values have been adjusted using the Tukey method for multiple comparisons, ≤ 0.05 are highlighted in bold. Table Key: est. = estimate, *SE* = standard error; *df* = degree of freedom. A = *Acopora*, Pv = *Pavona*, Pt = *Platygyra*.

### 3.3. Bioindicator Surveys

The total abundance of fish (Figure 6; Table 3) differed significantly among the sites (Table 4; Kruskal-Wallis χ*²* (2) = 54.1, *p* < 0.001, η*^2^* = 0.104). Post-hoc pairwise comparisons showed that the average fish abundance at the unrestored site (0.7 ± 1.1 *SD*) was significantly lower than both the reference (4.3 ± 4.9; *p* < 0.001) and the restored (5.1 ± 6.3; *p* < 0.001) sites. No significant difference was found between restored and reference sites (*p* = 1.0). The total abundance of invertebrates (Figure 6; Table 3) also differed significantly among the sites (Kruskal-Wallis χ*²* (2) = 7.33, *p* < 0.05, η*^2^* = 0.014). Post-hoc pairwise comparisons showed that the average invertebrate abundance was only different between the restored (77.7 ± 74.5) and the unrestored sites (47.1 ± 47.1, *p* < 0.05).

**Figure 6.**
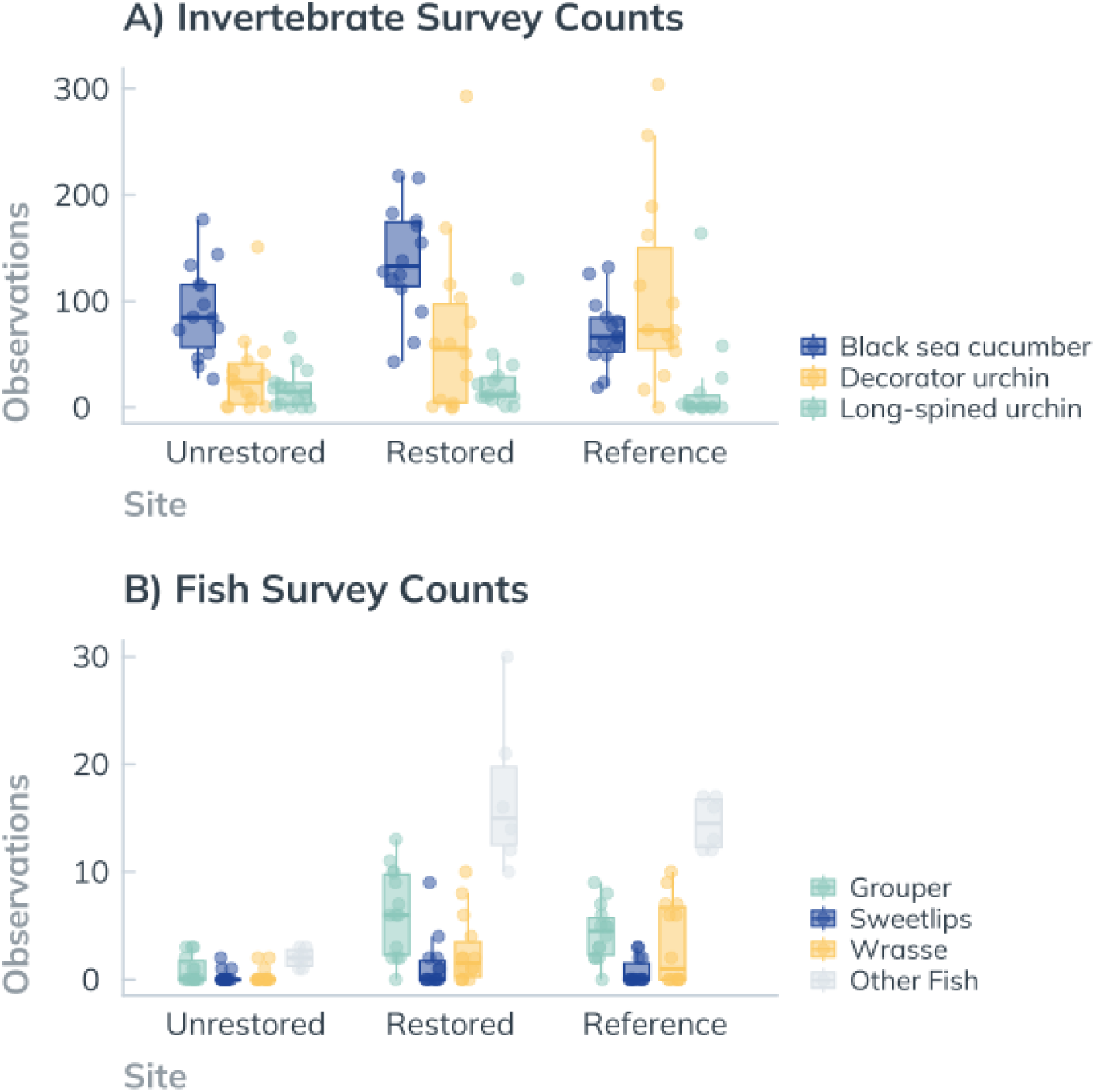
Visual survey counts. Each point represents the total number of observations of each bioindicator recorded during a single monitoring event for: A) Invertebrates and B) Fish.

**Table 3.** Total abundance of bioindicator sightings across 14 surveys, and the mean count and standard deviation (SD) for each survey at the reference, restored, and unrestored sites.

| Site |  | Reference |  |  | Restored |  |  | Unrestored |  |  |
| --- | --- | --- | --- | --- | --- | --- | --- | --- | --- | --- |
| Bioindicator | Type | Total count | Mean count | SD | Total count | Mean count | SD | Total count | Mean count | SD |
| Grouper | Fish | 61 | 4.4 | 2.6 | 87 | 6.2 | 4.0 | 13 | 0.9 | 1.3 |
| Other Fish | Fish | 87 | 14.5 | 2.4 | 103 | 17.2 | 7.3 | 12 | 2.0 | 0.9 |
| Seahorse | Fish | 0 | 0.0 | 0.0 | 0 | 0.0 | 0.0 | 0 | 0.0 | 0.0 |
| Sweetlips | Fish | 10 | 0.7 | 1.2 | 19 | 1.4 | 2.5 | 4 | 0.3 | 0.6 |
| Wrasse | Fish | 47 | 3.4 | 3.9 | 37 | 2.6 | 3.2 | 5 | 0.4 | 0.7 |
| Black sea cucumber | Invert | 992 | 70.9 | 32.6 | 1937 | 138.4 | 52.5 | 1263 | 90.2 | 43.3 |
| Decorator urchin | Invert | 1498 | 107.0 | 90.0 | 974 | 69.6 | 82.2 | 447 | 31.9 | 39.8 |
| Long-spined urchin | Invert | 269 | 19.2 | 44.8 | 351 | 25.1 | 31.1 | 251 | 17.9 | 19.5 |

**Table 4.** Results of the Kruskal-Wallis test comparing total abundance of fish and invertebrates between sites. *p*-values ≤ 0.05 are highlighted in bold. *df* = degree of freedom. Ref = reference site, Res = restored site, Un = unrestored site.

| Bioindicator group | Test statistic | df | $p$-value | $p$-value (Pairwise) |
| --- | --- | --- | --- | --- |
| Fish | 54.1 | 2 | **< 0.0001** | Ref - Res: 1.0<br>Ref - Un: < 0.0001<br>Res - Un: < 0.0001 |
| Invertebrate | 7.33 | 2 | **0.0256** | Ref - Res: 0.364<br>Ref - Un: 0.752<br>Res - Un: 0.0209 |

### 3.4. Environmental DNA

#### 3.4.1 DNA Sequencing Output by Phyla

The dataset consisted of 3,981 ASVs and 1,350,202 total reads across all samples following sequence analyses and quality control filtering. A total of 35 eukaryotic phyla were detected and assigned across Chromista (9 phyla), Plantae (4 phyla), Animalia (14 phyla), Fungi (4 phyla), and Protista (4 phyla; Figure A2). Of these kingdoms, Chromista was represented by the greatest number of both ASVs and untransformed sequencing reads 2,544 ASVs; 812,859 reads) — followed by Plantae (541 ASVs; 98,190 reads), Animalia (397 ASVs; 393,712 reads), and Fungi (295 ASVs; 23,698 reads). There were 8 ASVs (381 reads) that could not be assigned to the kingdom level.

Of the five phyla with the highest sampled ASV richness, four were Chromista: Bacillariophyta (1,613 ASVs; 623,144 reads), Ochrophyta (389 ASVs; 67,893 reads), Ciliophora (218 ASVs; 76,469 reads), and Oomycota (202 ASVs; 23,437 reads; Figure 7). Rhodophyta however had the second-greatest assigned ASV richness of any phyla (457 ASVs; 64,298 reads), accounting for the majority (84%) of ASVs assigned to the kingdom Plantae (Figure 7). Among metazoans: Cnidaria had the greatest ASV richness (174 ASVs; 10,920 reads), followed by Mollusca (67ASVs; 167,474 reads), Porifera (46 ASVs; 3,016 reads), Platyhelminthes (32 ASVs; 7,184 reads), and Arthropoda (26 ASVs; 35,279 reads; Figure A2). In terms of raw sequencing reads, Annelida was the third most abundant phyla among all kingdoms while being represented by low ASV diversity (8 ASVs; 153,520 reads; Figure 7).

**Figure 7.**
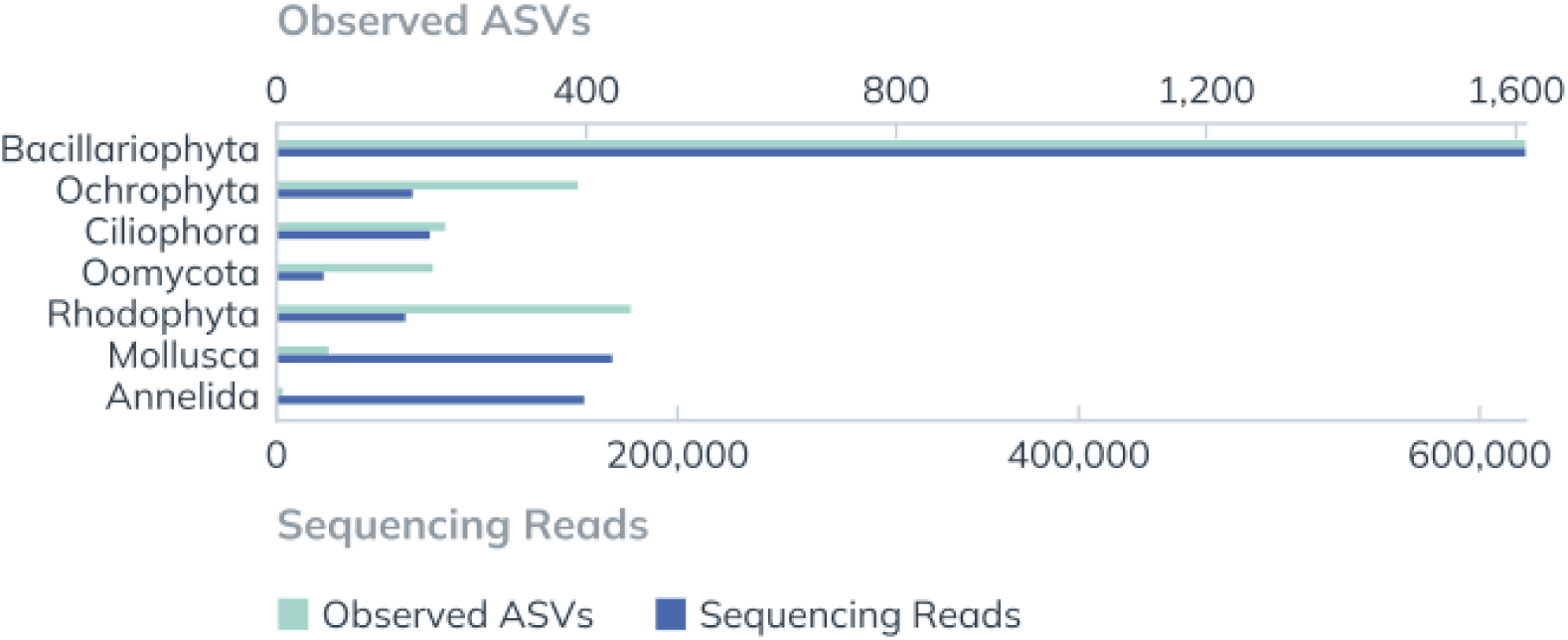
The number of observed ASVs and sequencing reads by phylum across all samples. Data shown reflects only those ASVs and reads which passed the prevalence and abundance filters prior to Hellinger transformation. For enhanced visibility, only the five phyla with the greatest ASV diversity (Bacillariophyta, Ochrophyta, Ciliophora, Oomycota, Rhodophyta) and/or the five phyla with the highest raw sequencing read counts (Bacillariophyta, Mollusca, Annelida, Ciliophora, Ochrophyta) were plotted.

#### 3.4.2. ASV Richness by Site

ASV richness was evaluated in terms of both the overall site richness and the average number of ASVs observed per sample (Figure 8). The restored site exhibited the highest overall richness, with 1,921 observed ASVs — 13.3% more ASVs than the number detected at the reference coral community (1,696 ASVs) and 23.5% more than the sampled richness of the unrestored seabed (1,555 ASVs). When excluding the Bacillariophyta ASVs — given the impact the disproportionately high diversity of this phylum may have on observed patterns of overall site richness — the restored site still displayed the highest number of observed ASVs (1,129). In contrast, a greater number of non-Bacillariophyta ASVs were observed at the unrestored site (1,003) than at the reference site (833). Non-Bacillariophyta richness at the restoration site was therefore 12.5% higher than that of the unrestored site and about 35.6% higher than that of the reference site.

**Figure 8.**
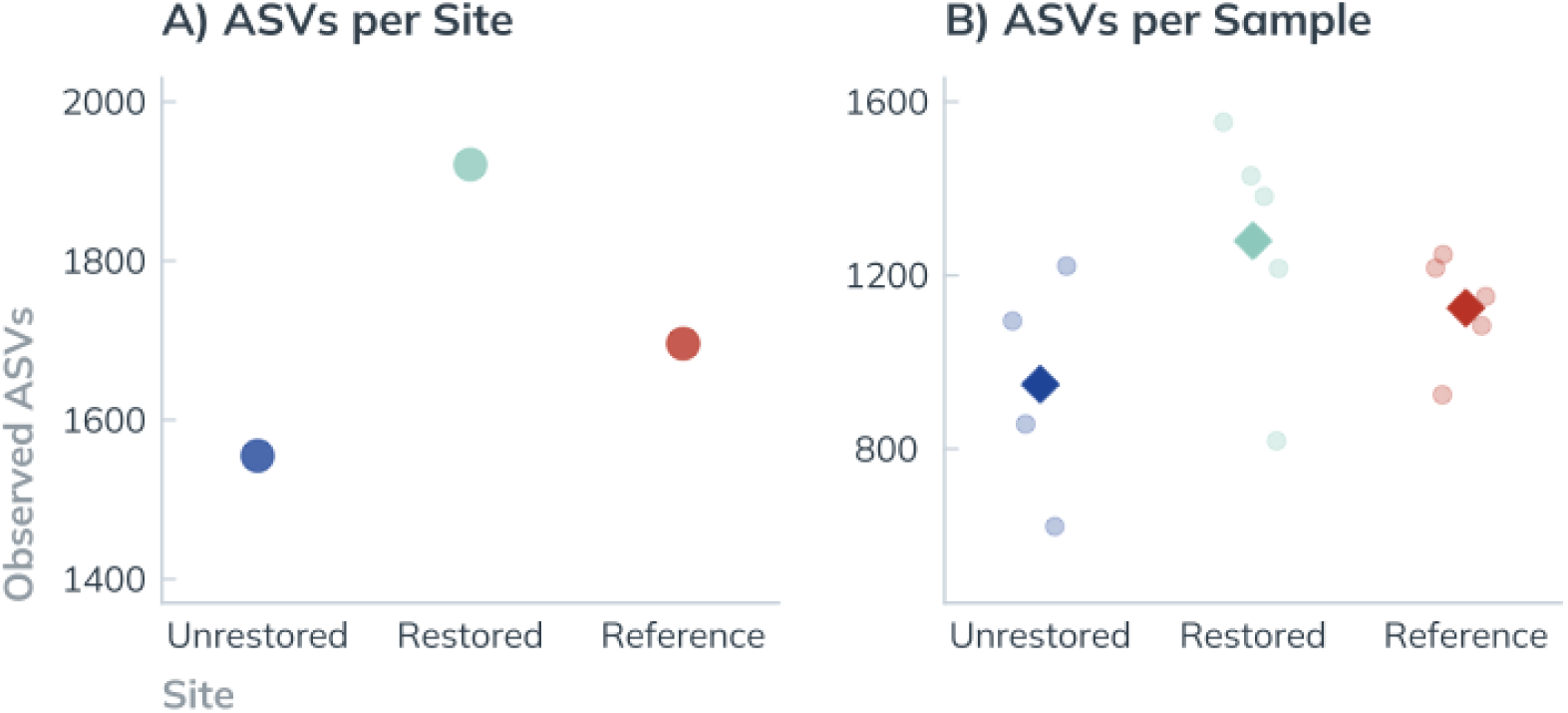
Point plots of observed ASV richness by site and per sample. Based on the parameters of the prevalence filter applied to the community metabarcoding dataset, all ASVs represented in this plot appeared in a minimum of three samples per site. (A) The number of unique ASVs observed across all samples from the unrestored (1,555 ASVs, *n* = 4), restored (1,921 ASVs, *n* = 5), and reference (1,696 ASVs, *n* = 5) site. Each ASV was counted once per site, regardless of how many replicates it was detected in. (B) The number of ASVs detected per sample are indicated by the circular points. Diamonds indicate mean ASV richness across all replicates for each site: 946 ± 266 ASVs per sample (±*SD*) from the unrestored (*n* = 4), 1,280 ± 284 from the restored (*n* = 5), and 1,125 ± 129 from the reference (*n* = 5) site.

Across all three sites, an average richness of 1,130 ASVs per sample was observed (± 255 *SD*, *n* =14). The restored site showed the highest average sample richness at 1,280 ASVs (± 284 *SD*, *n* = 5), followed by the reference site at 1,125 ASVs (± 129 *SD*, *n* = 5), and the unrestored site, which had the lowest average sample richness of 948 ASVs (± 266 *SD*, *n* = 4). Greater variation in ASV richness was observed among samples collected from the unrestored and restored sites than among those from the reference coral community (Figure 8B). Considering the relative differences in mean sample richness between sites, ASV richness from the restoration site was 13.7% greater than that of the reference community, and 35% higher than that of the unrestored site. However, the results of a one-way ANOVA of ASV richness by site indicate that the differences in sample ASV richness between sites were statistically insignificant (*F*(2, 11) = 2.225, *p* > 0.1; Table 5).

**Table 5:** The results of a one-way ANOVA testing for differences in ASV richness by site. . The ANOVA was run using the ‘vegan’ package for *R* (*(aov(Observed ASVs ∼ Site)*).

|  | Df | Sum Sq | Mean Sq | F value | Pr(>F) |
| --- | --- | --- | --- | --- | --- |
| **Site** | 2 | 243432 | 121716 | 2.225 | 0.154 |
| **Residuals** | 11 | 601744 | 54704 |  |  |

#### 3.4.3. Community Similarity

Most ASVs were unique to an individual site, as 76% of all ASVs were exclusively found among samples from one of the three locations (Table 6). A similar number of site-specific ASVs were found at the restoration site (1,059; 27% of all ASVs) and the reference site (1,064; 27%). In contrast, the unrestored site had a lower number of site-specific ASVs than the other two sites, with only 880 site-specific ASVs (22% of all ASVs). Among ASVs that were shared between only two sites, the restoration site shared a slightly higher number of ASVs with the nearby unrestored seabed (341 ASVs; 9% of all ASVs) than with the reference coral community (303 ASVs, 8% of all ASVs). However, the unrestored site had far fewer ASVs in common with the reference site (121 ASVs; 3% of all ASVs).

**Table 6:** Unique, ubiquitous, and shared ASVs by site.

|  |  | Observed ASVs | % of all ASVs | % of Reads<br>*Hellinger Transformed* | % of Reads<br>*No Transformation* |
| --- | --- | --- | --- | --- | --- |
| **Unique** | Unrestored | 880 | 22% | 11% | 18% |
|  | Restored | 1059 | 27% | 14% | 7% |
|  | Reference | 1064 | 27% | 19% | 13% |
| **Shared** | Unrestored -- Restored -- Reference | 213 | 5% | 23% | 27% |
|  | Unrestored -- Restored | 341 | 9% | 14% | 19% |
|  | Unrestored -- Reference | 121 | 3% | 5% | 6% |
|  | Restored -- Reference | 303 | 8% | 15% | 11% |
| **Total** |  | **3981** | **76%** |  |  |

Only 213 ASVs were identified across all three sites (Table 6). While these ubiquitous ASVs accounted for a small share (5%) of all observed ASVs, they were represented by a relatively large number of reads, accounting for 23% of Hellinger-transformed read counts across all samples. The majority of these ubiquitous ASVs were Bacillariophyta (123 ASVs; 57 of all ubiquitous ASVs), followed by Ochrophyta (21; 10%), Ciliophora (12; 6%), and Rhodophyta (11; 5%). All other phyla were represented by fewer than 10 ubiquitous ASVs. Two sets of Principal Coordinate Analyses (PCoA) were conducted to assess differences in community composition between sites. Each site was shown to have a distinct community structure, with both ordinations producing tight, largely non-overlapping clusters (Figure 9). The Jaccard distance ordination (presence/absence) showed some overlap between the unrestored and restored sites along the first principal coordinate (PCo1, 31.5%). However, all three sites occupied distinct positions along the second principal coordinate (PCo2, 24.3%; Figure 9A). PERMANOVA analyses of the Jaccard dissimilarity matrix confirmed significant differences in assemblages between the sites (*F* = 6.9, *p* < 0.001; Table 7), with post-hoc pairwise testing indicating significant differences between all site pairs (*p* < 0.01; Table 8).

**Figure 9.**
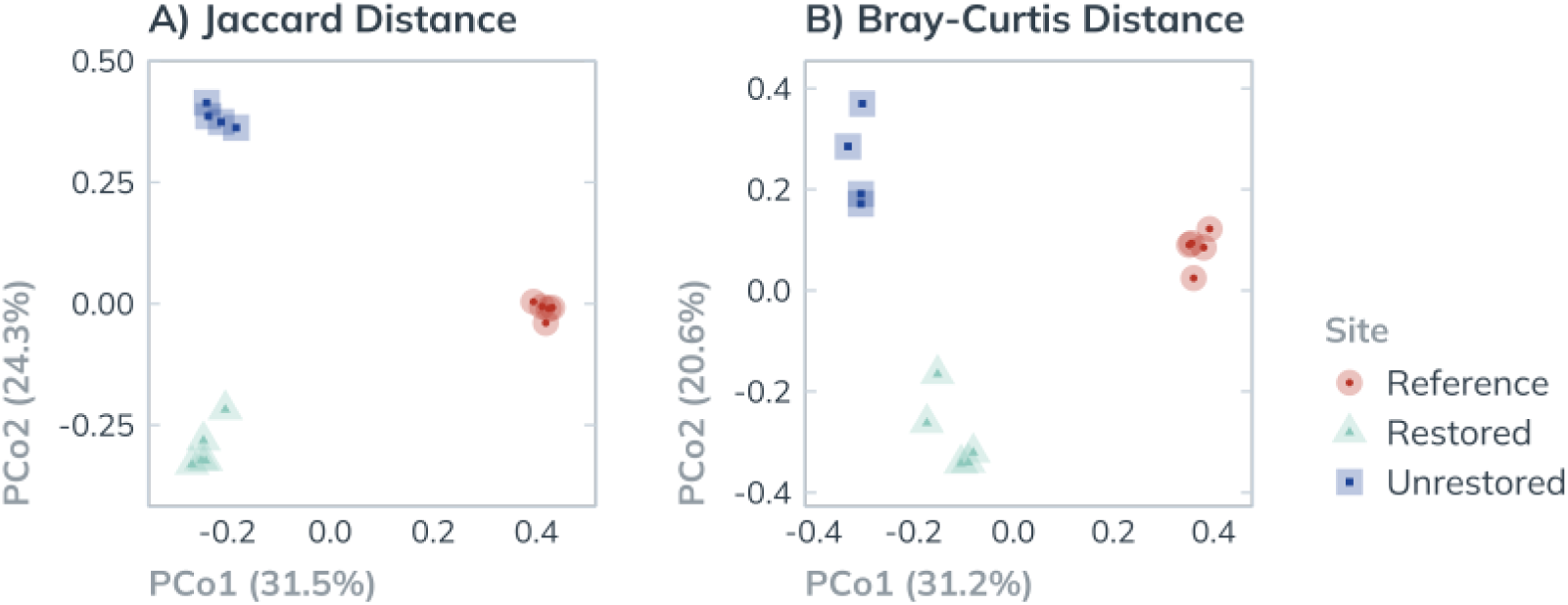
Beta diversity of eukaryotic ASVs. Dissimilarity in community composition was measured by community dissimilarity matrices. Each point represents a single sample, sites are distinguished by color and shape (A) PCoA ordination based on Jaccard distance, a presence-absence based metric. (B) PCoA based on Bray-Curtis distance, which incorporates abundance data (Hellinger-transformed).

**Table 7.** PERMANOVA results, testing for differences in eDNA community composition by site. Results were calculated separately from both a Jaccard Distance matrix and a Bray-Curtis Dissimilarity matrix using the ‘adonis2’ function of *vegan* (999 permutations).

|  |  | Df | SS | R<sup>2</sup> | F Model | p-Value |
| --- | --- | --- | --- | --- | --- | --- |
| Jaccard | Site | 2 | 2.377 | 0.555 | 6.859 | 0.001 |
|  | Residual | 11 | 1.906 | 0.445 |  |  |
|  | Total | 13 | 4.284 | 1.000 |  |  |
| Bray-Curtis | Site | 2 | 1.867 | 0.510 | 5.714 | 0.001 |
|  | Residual | 11 | 1.797 | 0.490 |  |  |
|  | Total | 13 | 3.664 | 1.000 |  |  |

**Table 8:** Pairwise Comparisons of beta diversity by site. Produced using the ‘pairwiseAdonis’ wrapper for ‘vegan’ (999 permutations) based on both Jaccard Distance and Bray-Curtis dissimilarity matrices.

|  |  | Restored – Reference |  |  |  |  | Restored – Unrestored |  |  |  |  | Reference – Unrestored |  |  |  |  |
| --- | --- | --- | --- | --- | --- | --- | --- | --- | --- | --- | --- | --- | --- | --- | --- | --- |
|  |  | Df | SS | R<sup>2</sup> | F | p | Df | SS | R<sup>2</sup> | F | p | Df | SS | R<sup>2</sup> | F | p |
| Jaccard | Site | 1 | 1.28 | 0.50 | 7.91 | 0.01 | 1 | 1.03 | 0.45 | 5.73 | 0.02 | 1 | 1.24 | 0.50 | 6.92 | 0.01 |
|  | Residual | 8 | 1.30 | 0.50 |  |  | 7 | 1.26 | 0.55 |  |  | 7 | 1.26 | 0.50 |  |  |
|  | Total | 9 | 2.58 | 1.00 |  |  | 8 | 2.29 | 1.00 |  |  | 8 | 2.50 | 1.00 |  |  |
| Bray-Curtis | Site | 1 | 0.95 | 0.45 | 6.42 | 0.01 | 1 | 0.76 | 0.38 | 4.23 | 0.01 | 1 | 1.10 | 0.49 | 6.60 | 0.01 |
|  | Residual | 8 | 1.18 | 0.55 |  |  | 7 | 1.25 | 0.62 |  |  | 7 | 1.16 | 0.51 |  |  |
|  | Total | 9 | 2.12 | 1.00 |  |  | 8 | 2.01 | 1.00 |  |  | 8 | 2.26 | 1.00 |  |  |

The Bray-Curtis distance analysis, which is calculated based on both occurrence and abundance, produced clear site clusters with minimal overlap. The restored site plotted between the unrestored and reference sites on the first principal component (PCo1, 31.2%), while the reference site again appeared centrally positioned between the other two on the second component (PCo2, 20.6%; Figure 9B). The PERMANOVA based on the Bray-Curtis dissimilarity matrix indicated statistically significant differences in community composition by site (*p* < 0.001; Table 7), with post-hoc tests confirming significant differences between all site pairs (*p* < 0.05 to 0.012; Table 8). In both ordinations, samples from the reference site clustered more closely together than those from the other two sites (Figure 9). This effect was particularly pronounced in the Bray-Curtis ordination, suggesting that read count representation of individual ASVs was more consistent across samples from the reference site.

#### 3.4.4. Taxonomic Composition by Site: Diversity and Abundance

Given the wide taxonomic breadth of the sequencing data, it was only feasible to compare eukaryote community composition at the phylum level. Taxonomic composition was first assessed using presence–absence transformed data, illustrating the number of ASVs observed within each phylum by site. ASV richness for each phylum was relatively consistent across the three sites, except for Bacillariophyta, which was represented by more ASVs and contributed a greater share of site richness at the reference site (Figure 10). Of the 36 phyla identified, 25 were represented by at least 10 ASVs and were retained to test for variation in phylum ASV richness by site (Figure A2). This test confirmed a significant difference in phylum richness by site (χ² = 138.1, *df* = 48, *p* < 0.001), primarily driven by differences in the number of Bacillariophyta observed between sites. Bacillariophyta ASV richness was highest at the reference site (863 ASVs, 50.9% of site ASVs), lower at the unrestored site (552 ASVs, 35.5%), and intermediate at the restored site (792 ASVs, 41.2%). Across all three sites, Bacillariophyta contributed 34.6% to the calculated chi-square value, while no other phylum contributed more than 6%. Overall, Bacillariophyta accounted for 40.5% of all detected ASVs. Recognizing that the high diversity of Bacillariophyta may obscure richness patterns among other phyla, a second chi-square test was performed excluding Bacillariophyta ASVs. In this analysis, ASV richness by phylum was largely equal across sites (χ² = 138.1, *df* = 46, *p* = 0.101). Cnidaria was the only potential exception, contributing 17.0% to the test statistic, and was slightly more diverse at the reference (79 ASVs, 9.5% of all non-Bacillariophyta ASVs) and restored (75 ASVs, 6.6%) sites than at the unrestored site (58 ASVs, 5.8%).

**Figure 10.**
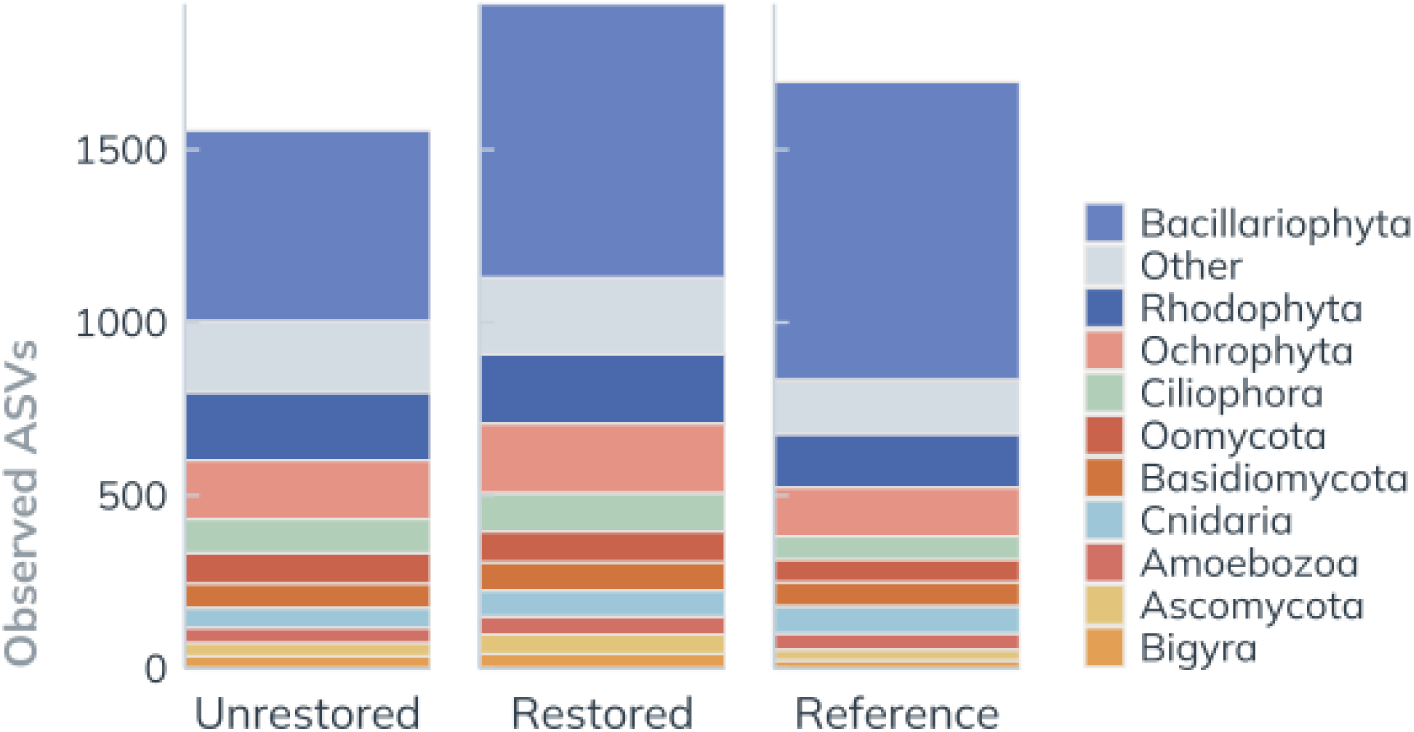
Eukaryotic phylum richness, by site. Each bar represents the total number of ASVs observed across samples from each site, separated by color to depict phylum-level ASV richness. Bars are colored to indicate the 10 most diverse phyla within the eDNA COI metabarcoding dataset. “Other” represents the sum of observed ASVs that belong to the remaining 15 phyla.

The taxonomic composition of both individual samples and merged site communities was further examined using Hellinger-transformed read abundances. Across all sites, Bacillariophyta was the most abundant phylum, accounting for an average of 51.1% of all transformed read counts per sample (± 11.8% *SD*; Figure 11A). The next two most abundant phyla, Ochrophyta and Rhodophyta, were generally present in similar proportions, each representing approximately 8% of total transformed read counts per sample (8.3 ± 1.4% and 8.0 ± 1.5% *SD*, respectively; Figure 11A). Phylum-level read abundance was more consistent among replicates from the reference community, mirroring the tighter clustering observed among reference site samples in the Bray-Curtis ordination (Figure 9B). In contrast, there was substantial variation in phylum representation across samples from the unrestored site. For the restored site, variation in phylum-level abundance was intermediate, with relative read abundance profiles for the seven most abundant phyla closely matching those of the reference site. However, two samples from the restored site had unusually high Mollusca read abundances (14.0% and 7.6% of total transformed reads).

**Figure 11.**
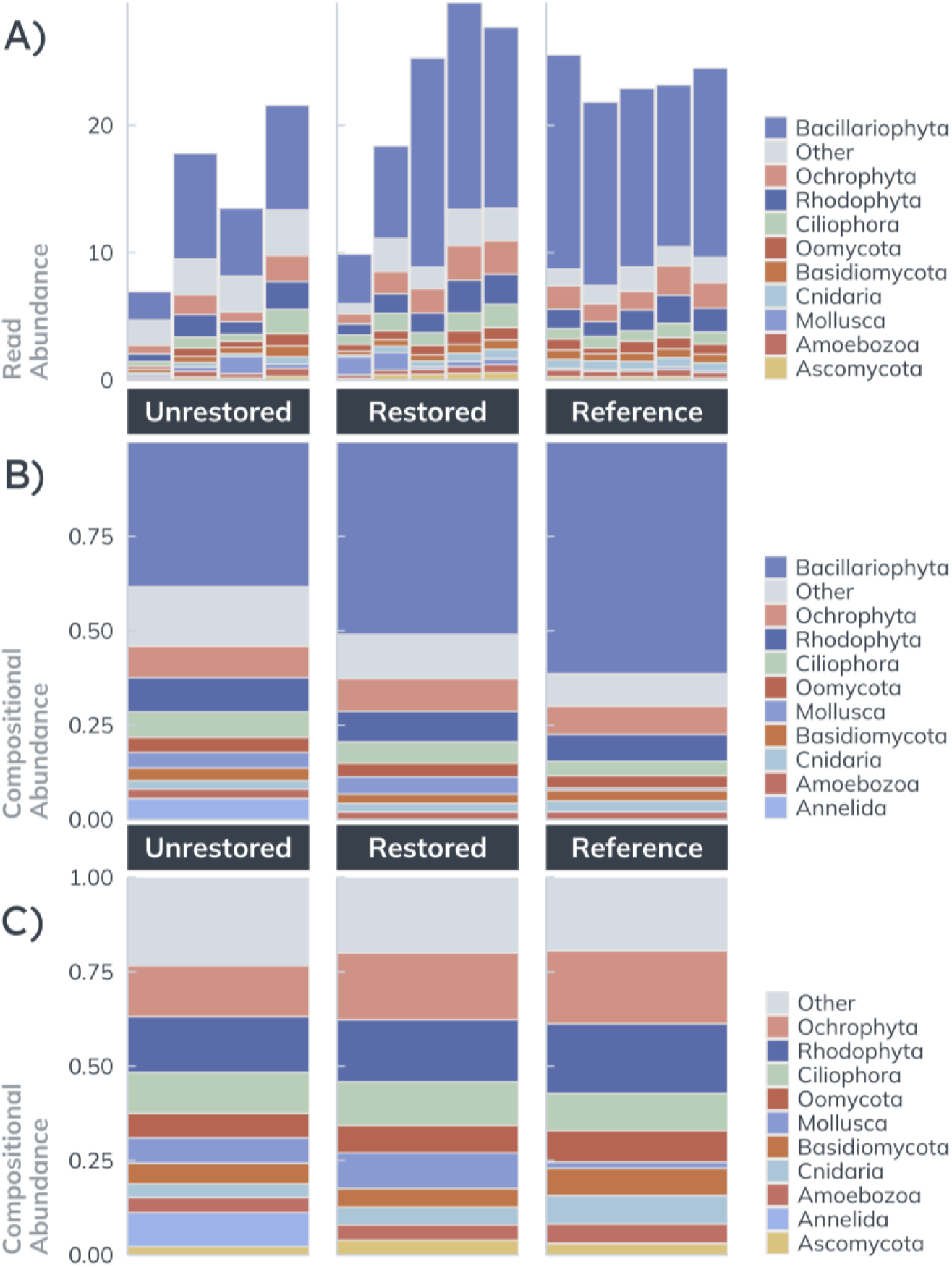
Taxonomic Composition by Phyla, based on Hellinger-Transformed Read Abundance. For each panel, the ten most abundant phyla are given distinct colors, while less abundant phyla have been merged as “Other”. Each phylum is depicted with the same color across all panels, but the ten most abundant phyla vary by panel as a result of differences in how data for each was processed (A) Stacked bar plot depicting the total Hellinger-transformed read abundance for each sample. (B) Read abundances were merged by site prior to Hellinger transformation, and subsequently converted to Compositional Abundance, to show the relative contribution of each phylum to the composite sequenced community for each site. (C) Bacillariophyta ASVs were removed prior to merging by site and Hellinger transformation, to more clearly depict differences in the proportions of non-Bacillariophyta reads assigned to other phyla between sites.

Compositional abundance—presented as relative Hellinger-transformed abundance by site—was used as a proxy for site-level community composition. Bacillariophyta abundance varied considerably by site, with the highest compositional abundance at the reference site (61.4%), followed by the restored (50.9%) and unrestored (38.4%) sites (Figure 11B), consistent with the gradient in Bacillariophyta richness observed across sites (Figure 10). As Bacillariophyta accounted for 53.7% of all transformed read counts across samples, patterns in compositional abundance for other phyla were obscured, prompting removal of Bacillariophyta from the dataset to better examine differences among remaining phyla.

Following Bacillariophyta, Ochrophyta and Rhodophyta were the most abundant phyla across all sites. For both algal phyla, compositional abundance was lowest at the unrestored site, intermediate at the restored site, and highest at the reference site. In samples from the unrestored, restored, and reference sites, respectively: Ochrophyta accounted for 13.5%, 17.7%, and 19.3%, and Rhodophyta for 14.8%, 16.5%, and 18.4% of non-Bacillariophyta transformed reads (Figure 11C). A similar gradient was observed for relative Cnidaria abundance, which was lowest at the unrestored site (3.6%), higher at the restored site (4.6%), and highest at the reference site (7.5%; Figure 11C).

Other phyla displayed different patterns of compositional abundance between sites. Ciliophora abundance was relatively even across all samples, being slightly higher at the unrestored (10.8%) and restored (11.5%) sites than at the reference site (9.9%; Figure 11C). In contrast, Basidiomycota was more abundant at the reference site (7.1%) than at the unrestored (5.5%) and restored (5.0%) sites. Differences in relative Mollusca sequence abundance were more pronounced, being high at the unrestored (6.7%) and restored (9.5%) sites, but negligible at the reference site (1.7%). Notably, Annelida accounted for a large share of transformed reads at the unrestored site (9.0%) but contributed minimally to the read counts of the other two sites (<1.0%). Ascomycota sequences accounted for a greater share of reads from the restored site (3.9%) than from the reference (2.9%) or unrestored (2.2%) sites.

## 4. Discussion

Coral restoration has two primary ecological goals, broadly: coral population enhancement, and the preservation of community biodiversity and ecosystem function. In this study, we deployed a novel 3D-printed ceramic tile design to establish a coral community in a marine park in Hong Kong. Evaluation of the tiles’ efficacy for coral reef restoration showed that the artificial structures provided a suitable substrate, supporting sustained coral growth and high survivorship of transplanted fragments. This, in turn, resulted in a measurable enhancement of site biodiversity.

### 4.1. Ecological Outcomes

#### 4.1.1. Coral Performance

Our results demonstrate that the 3D-printed ceramic tiles provided a suitable substrate for coral attachment and growth. After four years, 91% of the 378 coral fragments outplanted to the restoration site remained alive, with only 2% showing signs of tissue loss. This survivorship rate significantly exceeded the average of 66% reported for coral transplantation projects — a figure often considered an overestimate, as most studies monitor survivorship for only up to 18 months post-transplantation [11,12]. Our project met the suggested targets of the Coral Restoration Consortium (CRC) of the National Oceanic and Atmospheric Administration (NOAA), whose Restoration Evaluation Tool provides the only comprehensive suite of quantitative targets for evaluating coral reef restoration performance [49].

The CRC recommends the following criteria for coral health and survivorship, measured one year after transplantation: (1) high coral survivorship (>80% of outplanted fragments alive), (2) high mean live tissue coverage (>80% per colony), (3) limited coral bleaching (<5% of fragments with tissue loss due to bleaching), and (4) low disease prevalence (<10% of fragments showing disease)[49]. At the one-year mark, all criteria were met: only one fragment died, and five detached, resulting in an overall survivorship rate of 98%. While the exact percentage of tissue loss per fragment was not quantified, 98% of fragments exhibited full tissue cover, suggesting mean live tissue coverage exceeded 80%. Diver surveys using color indices found no signs of coral bleaching or disease. The CRC’s longer-term targets for survivorship are >65% cover for years 2–5 post-transplantation and >50% thereafter. Although our monitoring covers only the first four years, the high final survivorship and minimal losses over time suggest the project will continue to meet these criteria, barring unforeseen acute stress events.

Coral growth rates can vary considerably among species and morphologies, with branching corals generally growing faster than massive and plating forms [50,51]. Consistent with expectations, we observed the fastest growth in *Acropora* (0.35 cm m^-1^), followed by *Pavona* (0.21 cm m^-1^) and *Platygyra* (0.15 cm m^-1^) over four years. These growth rates are consistent with our expectations, based on their respective morphologies, as branching forms typically show faster growth due to their structural adaptations that optimize light capture and nutrient acquisition [52]. It is, however, difficult to contextualize the measured growth rates for this project relative to other studies, as environmental conditions, such as temperature, light, water quality, nutrient availability, salinity, and aragonite saturation also affect coral growth [53,54]. As what can be considered a “successful” growth rate is highly context-specific, the Coral Restoration Consortium (CRC) does not specify specific targets for coral growth rates [49]. Rather, it considers one successful outcome of outplanting to be an enhancement of coral reef structure and complexity, measured as an increase in mean coral height or linear extension [49]. Our project satisfied this criterion, with the mean maximum linear extension of all three genera, at minimum, doubling.

*Acropora* are often favored for restoration projects, as their relatively high growth rates can support rapid establishment, while their branching morphology enhances structural complexity [55,56]. In our study, *Acropora* achieved the highest survivorship, albeit with a higher incidence of breakage compared to *Pavona* and *Platygyra*. The higher incidence of breakage among transplanted *Acropora* fragments is consistent with the ecological characteristics of corals with branching morphologies, which reproduce via fragmentation and employ rapid growth to overcome the high-disturbance environments of coral reefs [57,58]. However, the high survivorship of *Acropora* observed here is somewhat atypical; Within coral restoration literature, it is often acknowledged that there is a tradeoff between growth and survivorship rates in transplanted *Acropora* fragments, which often exhibit much higher mortality rates than other slower-growing morphospecies [49,59–61].

Transplanting diverse coral assemblages offers ecological advantages, as fast-growing branching species like *Acropora* can offset predation and disturbance pressures on slower-growing taxa, facilitating their establishment [62]. Massive and foliose corals tend to be more resistant to environmental stressors, contributing to long-term stability [62]. Functional diversity also enhances community-level resilience, with mixed assemblages of autotrophic and heterotrophic species conferring increased resistance to thermal bleaching [63]. This approach is especially relevant in urbanized, eutrophic environments such as Hong Kong, where heterotrophic taxa may be more resilient under nutrient enrichment [28,64].

Despite these benefits, implementing species-diverse outplanting strategies presents challenges, particularly in optimizing attachment techniques for different morphologies. In our study, most fragment losses resulted from detachment, with *Pavona* (a plate coral) being disproportionately affected. This outcome highlights the need to further develop attachment methods tailored to specific growth forms to maximize survivorship and overall restoration efficacy. Addressing these technical considerations will enhance the ecological benefits of species and functional diversity, increasing the resilience and sustainability of restored communities. In summary, our monitoring demonstrates that 3D-printed ceramic tiles provide a suitable substrate for coral attachment and growth. However, it is crucial to recognize that while the substrate plays a significant role in this success, appropriate site selection ultimately determines coral survivorship outcomes.

#### 4.1.2. Biodiversity

##### 4.1.2.1. Fish and Macroinvertebrate Abundance

One of the principal considerations in designing the reef tiles was to create an artificial structure that would attract and support a diverse community of marine taxa. Given the high economic and nutritional value of fish to coastal communities, fish have long been the central — and often sole — focus of studies on how coral-associated organisms are affected by the construction of artificial reefs [12,65]. A number of studies have demonstrated that fish are attracted to artificial structures, which are commonly associated with more abundant and diverse fish assemblages [66–69]. In this study, visual surveys found an average of seven times more fish at the restoration site than at the nearby unrestored seabed, with abundances comparable to those at the natural reference coral community. Macroinvertebrate abundance similarly differed between sites, with 45% greater abundance of sea urchins and sea cucumbers at the restored site than at the unrestored area.

Moderate densities of sea urchins and small herbivorous fishes on and near the tiles indicate promising progress toward establishing a functional coral ecosystem. Herbivory plays an essential role in maintaining coral cover and diverse benthic assemblages — particularly in eutrophic environments like Hong Kong, where high nutrient levels favor the proliferation of fast-growing algae that can outcompete corals for benthic cover and reduce biodiversity [70–72]. While moderate sea urchin densities help control macroalgae, excessive urchin densities can harm corals [73,74]. Our surveys found comparable urchin densities at the restoration and reference sites, which suggests the urchin abundance noted at the restoration site is compatible with coral growth. Looking beyond herbivory, the surveyed fish taxa included representatives of different trophic levels — sweetlips (Plectorhinchus spp.) that feed on shrimp and small crabs, wrasse (Labridae) which prey on larger invertebrates such as sea urchins and mollusks, and groupers (Epinephelinae) known to feed on other reef fish, octopuses, and larger crustaceans. The presence of fish reliant on a diversity of feeding strategies suggests that the restored habitat is supporting a functioning trophic network. From a socioeconomic perspective, the presence of commercially valuable species such as groupers also enhances the potential for local community engagement and economic viability of restoration efforts. Our results meet the CRC’s criterion for non-coral community biodiversity enhancement, which requires greater abundances of fish and coral grazers at the restoration site relative to a control or pre-restoration baseline.

The increased abundance of fish and macroinvertebrates at the restoration site may reflect either true enhancement of local carrying capacity or merely an aggregation around the new habitat, a distinction particularly relevant for highly mobile species like fish [11,67]. Without pre-restoration biodiversity data, it is unclear whether elevated abundances result from local recruitment and population growth or redistribution from adjacent areas. However, having multiple species aggregate, even if they are of similar functional groups, provides functional redundancy — which enhances both ecosystem function and resilience [75]. While visual surveys are effective for monitoring large, mobile taxa, they often underrepresent cryptic or small-bodied species [76]. Future assessments should incorporate pre-installation baselines and complementary methods, such as environmental DNA, to better evaluate biodiversity and carrying capacity impacts.

##### 4.1.2.2. eDNA Cryptobiome

The rugose tile surfaces, combined with the growth of transplanted coral fragments, served to enhance the structural complexity and resource availability of a previously uniform sandy seabed. Such complexity is widely recognized as a key driver of both macro- and microfaunal diversity in reef systems [67,69]. Indeed, the restored site exhibited 23.5% greater eukaryotic richness (1,921 ASVs) compared to that of the unrestored seabed (1,555 ASVs). Additionally, more unique ASVs were found at the restoration site than at the unrestored seabed (1,059 and 880 ASVs, respectively). Beta diversity analyses revealed distinct, non-overlapping clusters for each site, with PERMANOVA confirming significant differences in community composition between sites (*p* < 0.001). Our findings indicate that, despite the close proximity of the two sites (50 m), the restoration project produced a more diverse, distinct community assemblage from that of the unrestored seabed. Ecological theory generally posits that elevated taxonomic richness enhances ecosystem resilience and functional stability — even if much of the diversity is functionally redundant [77–79]. Under this theory, the increased richness associated with the restoration project would serve to enhance ecosystem resilience of the marine park.

Frameworks for ecosystem restoration are commonly framed around the goal of returning the community structure of a degraded ecosystem to that of an undisturbed baseline [80,81]. It is recommended that such objectives be evaluated through comparison to a reference site [81]. In our study, evidence for convergence between the eDNA communities sampled at the restoration and reference sites — such as comparable richness and increased read abundances of key taxa — was tempered by findings that the restored and unrestored sites continued to share certain community characteristics distinct from the natural reference coral community. Richness at the restored site (1,921 ASVs) exceeded that of the reference reef (1,696 ASVs). In terms of taxonomic composition: Both presence-absence and abundance data revealed gradients in the read abundance and ASV diversity of foundational groups such as Bacillariophyta, Rhodophyta, Ochrophyta, and Cnidaria between sites, which were greatest at the reference site, intermediate at the restored site, and least prevalent at the unrestored site. Artificial reef communities are known to undergo dynamic changes over years [82], with ecological succession continuing for years following deployment [83]. The differences in phylum-level community structure between the restoration and reference sites could also be partially attributed to having only sampled one point in time, two years after deployment, which captured only a snapshot of ongoing transition at the restoration site. While our results suggest that the community at the restoration site is becoming more similar to that of the reference community, these two sites are likely to retain distinct assemblages over the long term. Given the highly heterogeneous nature of urbanized marine ecosystems, variations in environmental conditions between the two sites will continue to shape each community in different ways [84,85]. Differences in substrate will also continue to influence the ongoing recruitment of benthic organisms to the site, as settlement preferences for different taxa vary based on substrate material and structure [86]. Rather than representing an incomplete transition to the reference community, the restoration site is likely developing into a distinct assemblage, which confers its own benefits to local biodiversity: Increased heterogeneity in both habitat and community structure between sites may ecosystem resilience by supporting a broader range of species and functional roles, improving the system’s capacity to respond to disturbances [87,88]. The addition of novel surfaces and substrates, combined with coral outplanting, can augment the seascape by introducing new ecological niches, facilitating coral community assemblages that are distinct from those found on natural reef ecosystems [89,90]. Coral restoration projects are typically implemented to enhance existing, though often degraded, coral communities — leveraging the knowledge that environmental conditions at the chosen restoration site are already generally suitable for coral growth. However, the results of this study highlight the potential benefits of installing artificial reefs in areas where all necessary conditions for coral growth met, aside from suitable substrate. The eDNA results demonstrate that it is possible to foster communities that are richer than those observed in neighboring natural reefs. It is, however, important to consider this finding within two specific contexts.

First, our reference coral community is located in a region that has experienced considerable degradation; thus, the richness observed at the restoration site was interpreted relative to a reasonable target for biodiversity given the highly urbanized environment. A similar pattern may not hold for restoration work conducted among less impacted reefs. Alternatively, the high richness sampled from the restoration site may have been facilitated by the relatively high regional biodiversity of Hong Kong [91]. Artificial reefs out planted in areas with less regional biodiversity to recruit from may be less successful at developing diverse communities. Second, unavoidable biases are introduced at every stage of eDNA-based monitoring — from differential preservation of DNA in the environment between taxa, to sampling, laboratory procedures, and bioinformatics. These biases can favor the detection of certain groups. While we observed increased overall richness, this measure, as with all survey methods, serves as a proxy for true site diversity. It is possible that richness gains reflect increases in specific taxa, while other groups may continue to be better represented at the reference site.

To our knowledge, this study represents the first application of eDNA to assess the outcomes of a coral reef restoration project. With no prior studies using sedimentary eDNA to compare marine communities following ecosystem restoration—or even across anthropogenic impact gradients—the ecological significance of the observed differences in richness and composition remains challenging to contextualize. While differences in richness between sites were not statistically significant, the effect size may still be ecologically relevant, particularly given the low replication in this study. As the temporal and spatial variability of eDNA in marine sediments remain poorly characterized [24], our findings are further limited by having only sampled a single time point. We recommend that future eDNA monitoring projects incorporate pre-deployment baseline sampling and annual monitoring to better capture the trajectory of succession and the persistence of these patterns.

While there are no relevant points of comparison in marine restoration ecology: In a comprehensive meta-analysis of over 400 papers, terrestrial restoration sites were found to exhibit, on average, 20% higher biodiversity than unrestored sites (across a range of indices, not eDNA specific)[92]. The 23.5% difference in richness we observed between the unrestored and restored sites is consistent with the effect of restoration in terrestrial environments. Although these findings cannot be directly equated due to differences in methodological approaches and ecosystem contexts, the parallel observed here is noteworthy and may indicate a broader pattern across restoration efforts.

### 4.2. Measuring Coral Restoration Outcomes

The most stated objective among peer-reviewed studies of coral reef restoration, whether conducted on existing reefs or in areas augmented with artificial structures, is the establishment of a self-sustaining, functioning coral reef ecosystem [21]. While the majority of coral restoration studies state objectives with a broad ecological scope — to preserve critical ecosystem functions, enhance biodiversity, and foster resilience — a review by Boström-Einarsson et al. found that 45% of restoration projects that noted ecological objectives failed to collect relevant measurements to evaluate them [11]. Across the published literature, there is an apparent misalignment between the ecological aspirations of restoration work and the indicators generally chosen to measure success, with fragment-scale measurements of coral growth and survivorship being reported more than twice as often than community-level species composition metrics [21,93]. Among reef restoration studies that include surveys of coral-associated communities, fish are often the sole taxa reported (∼60%)[12]. Despite eDNA having the potential to sample biodiversity across the tree of life, recent work employing eDNA analyses as a tool for monitoring reef restoration outcomes also suffers this bias: With one exception, all previous work using eDNA to characterize communities associated with artificial reefs and coral restoration have focused on fish populations [94–98].

Levy et al. published the first example of eDNA being used to detect a broad range of metazoan taxa settling on or near artificial structures in a marine environment, demonstrating the promise of eDNA for capturing the often-overlooked diversity of reef-associated organisms [99]. Building on this work, our study is the first to use eDNA metabarcoding to assess how coral reef restoration influences eukaryote diversity. By including both restored and control sites, our approach enables a direct comparison of biodiversity outcomes attributable to restoration interventions—an advancement over previous studies, which typically report the community that has settled on new structures without baseline or control site assemblages for comparison. To our knowledge, the work of Knoester et al. is the only benthic biodiversity survey to compare communities associated with artificial reefs to those of a control site; their visual surveys of several macroinvertebrate groups revealed different effects on abundance but could not resolve an overarching effect on site-wide biodiversity [100]. Our survey therefore represents the first evidence of a community-wide enhancement of site richness associated with a coral restoration project.

A major strength of using eDNA for coral ecosystem monitoring is its ability to detect a broad suite of taxa, including cryptic and sessile organisms such as crustose coralline algae (CCA) and sponges [99]. These groups are increasingly recognized for their beneficial roles in coral recruitment, ecosystem engineering, and overall reef resilience [101,102]. By capturing this wider spectrum of biodiversity, eDNA offers a more holistic perspective on the ecological outcomes of restoration interventions. However, several limitations must be acknowledged. The replication and temporal resolution of our eDNA sampling were constrained, limiting our ability to assess temporal trends or establish causality. More broadly, the novelty of eDNA applications in reef restoration means that there are few comparable datasets, complicating interpretation and benchmarking of restoration success. Additionally, questions remain about the ecological significance of eDNA detections, as the method can recover signals from transient or low-abundance taxa whose functional roles within the ecosystem are unclear [103].

Despite these challenges, eDNA provides a valuable foundation for developing more nuanced functional indices. While our study focused on biodiversity as an endpoint, eDNA data could also be leveraged to infer aspects of trophic complexity—an attribute increasingly recognized as central to ecosystem functioning and resilience [80]. Moreover, eDNA can inform on the presence and relative abundance of foundational versus non-native species, and track functional groups relevant to nutrient cycling, reproductive outputs, and habitat stratification. Realizing the full potential of these molecular tools will require further experimental validation and integration with other monitoring approaches, but our findings demonstrate the capacity of eDNA to advance the field toward more comprehensive and ecologically relevant assessments of coral reef restoration outcomes.

Our monitoring efforts demonstrate that the restoration project satisfied, and indeed exceeded, all relevant CRC evaluation criteria and restoration objectives. The 3D-printed ceramic tiles provided a stable substrate for coral survivorship and growth and supported higher abundances of key fish and macroinvertebrates. Furthermore, we demonstrate that sedimentary eDNA analyses can be used to effectively detect previously unmeasured changes in sitewide biodiversity, finding evidence that the tile deployment and coral transplantation supported increased eukaryote biodiversity at the restoration site. These results suggest that the restoration project was successful and support the potential for future use of these tiles as artificial substrates for coral restoration.

## Author Contributions

V.Y. and D.M.B. conceived the project and secured research funding. Monitoring surveys were organized and led by H.L., Z.W., P.T., and V.Y., with survey data processed by H.L. and Z.W. Coral growth and survivorship data analyses were conducted by J.W., supported by A.C., V.Y., S.M.B., and D.M.B. Supervision of eDNA sampling and laboratory work was provided by V.Y., with advice from S.M.B. and A.C. eDNA data analyses were performed by A.C., supported by S.M.B. and D.M.B. All data figures were created by A.C. Manuscript writing was led by A.C., with contributions from J.W. and V.Y. The manuscript was edited by all authors. All authors have read and agreed to the published version of the manuscript.

## Institutional Review Board Statement

Not applicable.

## Informed Consent Statement

Not applicable.

## Data Availability Statement

The data presented in this study are available on request from the corresponding author to comply with government contract terms.

## Acknowledgments

The authors gratefully acknowledge the Agriculture, Fisheries and Conservation Department (AFCD) for funding the project and providing permitting support essential to the implementation of this study. The authors also wish to thank the World Wide Fund for Nature, Hong Kong (WWF-HK) for their valuable logistical assistance. Jordan Pierce is acknowledged for his contributions to the early-stage design development of the 3D printed ceramic structure. Appreciation is extended to Professor Chris Webster, Christian Lange, Lidia Ratoi, and Dominic Co from the School of Architecture at the University of Hong Kong for their assistance in the fabrication of the prototypes. We also thank Haze Chung for her assistance processing the eDNA samples, Rainbow Tsang for her contribution to the eDNA bioinformatics, as well as all volunteers who helped with the deployment of the ceramic reef tiles in Hoi Ha Wan and subsequent surveys.

## Abbreviations

The following abbreviations are used in this manuscript:

HHWMP: Hoi Ha Wan Marine Park
eDNA: Environmental DNA

## Appendix A

**Table A1.** Comparison of generalized linear mixed models (GLMMs) for coral extension rates across genera, evaluated using AIC. K denotes the number of model parameters, logLik is the log-likelihood of the model, ΔAIC represents the difference in AIC relative to the best-fitting model, and AIC Weight indicates the relative likelihood of each model given the data. The best-fitting model is highlighted in bold.

| Model | K | logLik | AIC | ΔAIC | AIC Weight |
| --- | --- | --- | --- | --- | --- |
| Random: unit_no<br>AR1: survey_no | 7 | 18080.95 | -36147.91 | 168.75 | <0.001 |
| Random: unit_no/tile_no<br>AR1: survey_no | 8 | 18102.02 | -36188.05 | 128.61 | 0.003 |
| Random:<br>unit_no/tile_no/coral_no<br>AR1: survey_no | 9 | 18167.33 | -36316.66 | 0.00 | 0.997 |

**Table A2.** Cumulative number of the coral condition by genus (Acropora, Pavona, and Platygyra) during each monitoring survey. Status indicators include: H (healthy), PM (partial mortality), D (dead), and DTCH (detached).

|  | *Acropora* |  |  |  | *Pavona* |  |  |  | *Platygyra* |  |  |  |
| --- | --- | --- | --- | --- | --- | --- | --- | --- | --- | --- | --- | --- |
| *Survey* | H | PM | D | DTCH | H | PM | D | DTCH | H | PM | D | DTCH |
| 1 | 126 | 0 | 0 | 0 | 126 | 0 | 0 | 0 | 126 | 0 | 0 | 0 |
| 2 | 126 | 0 | 0 | 0 | 125 | 0 | 0 | 1 | 126 | 0 | 0 | 0 |
| 3 | 126 | 0 | 0 | 0 | 125 | 0 | 0 | 1 | 126 | 0 | 0 | 0 |
| 4 | 126 | 0 | 0 | 0 | 125 | 0 | 0 | 1 | 126 | 0 | 0 | 0 |
| 5 | 125 | 0 | 0 | 1 | 121 | 0 | 1 | 4 | 125 | 0 | 0 | 1 |
| 6 | 125 | 0 | 0 | 1 | 120 | 0 | 2 | 4 | 124 | 0 | 0 | 2 |
| 7 | 125 | 0 | 0 | 1 | 113 | 0 | 2 | 11 | 124 | 0 | 0 | 2 |
| 8 | 125 | 0 | 0 | 1 | 107 | 0 | 4 | 15 | 123 | 0 | 1 | 2 |
| 9 | 125 | 0 | 0 | 1 | 105 | 0 | 4 | 17 | 123 | 0 | 1 | 2 |
| 10 | 124 | 0 | 0 | 2 | 105 | 0 | 4 | 17 | 122 | 1 | 1 | 2 |
| 11 | 122 | 0 | 0 | 4 | 101 | 0 | 4 | 21 | 116 | 4 | 3 | 3 |
| 12 | 120 | 0 | 0 | 6 | 99 | 0 | 4 | 23 | 109 | 11 | 3 | 3 |
| 13 | 119 | 1 | 0 | 6 | 99 | 0 | 4 | 23 | 116 | 3 | 4 | 3 |
| 14 | 118 | 1 | 0 | 7 | 96 | 0 | 4 | 26 | 112 | 6 | 4 | 4 |

## Appendix B Additional figures pertaining to the results of eDNA analyses.

**Figure A1.**
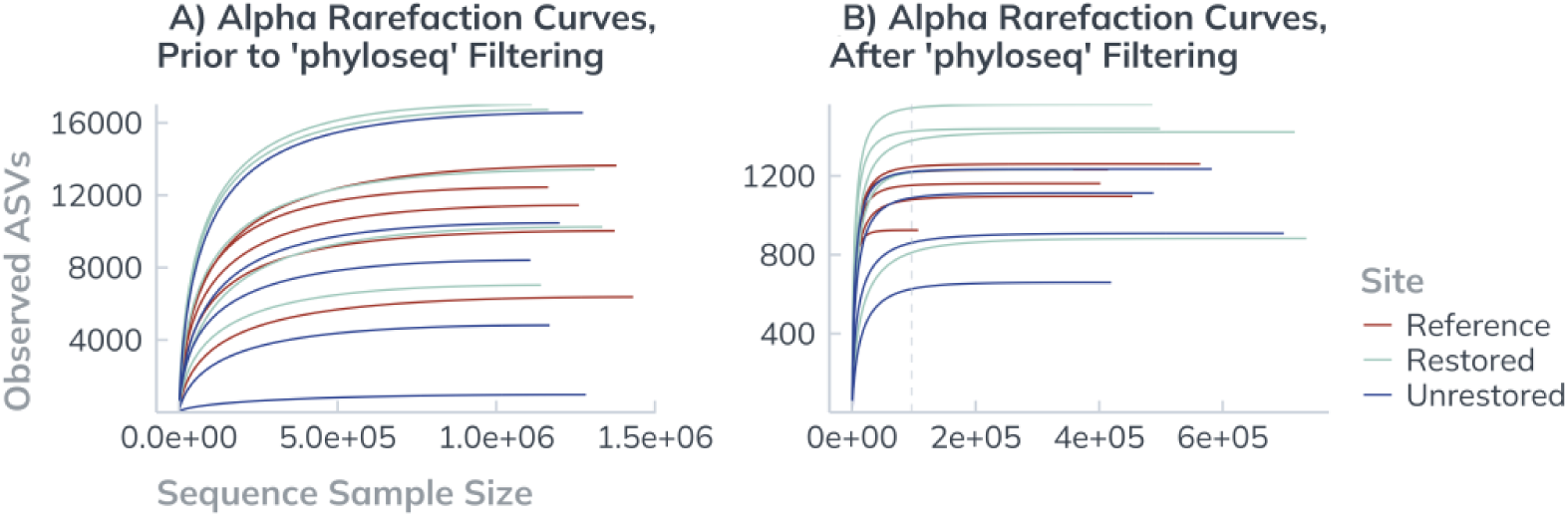
Rarefaction curves for ASV detection. Each line represents a sample, colored by site. (A) Rarefaction curves for ASVs inferred from the DADA2 *denoise-paired* function, before any subsequent cleanup steps. These curves indicate that the sequencing effort provided sufficient coverage for the taxa represented in the samples. The sample from the unrestored site that was subsequently excluded from analyses based on abnormally low richness (974 ASVs) and biased taxonomic composition is shown. (B) Rarefaction curves plotted from the phyloseq object that resulted following prevalence and abundance filtering, which retained only those ASVs that appeared in at least three samples from the same site and were represented by a minimum of 50 reads. The dotted grey line indicates the sampling depth used for rarefaction (96,443 reads), which was set to 90% of the depth of the sample with the fewest reads.

**Figure A2:**
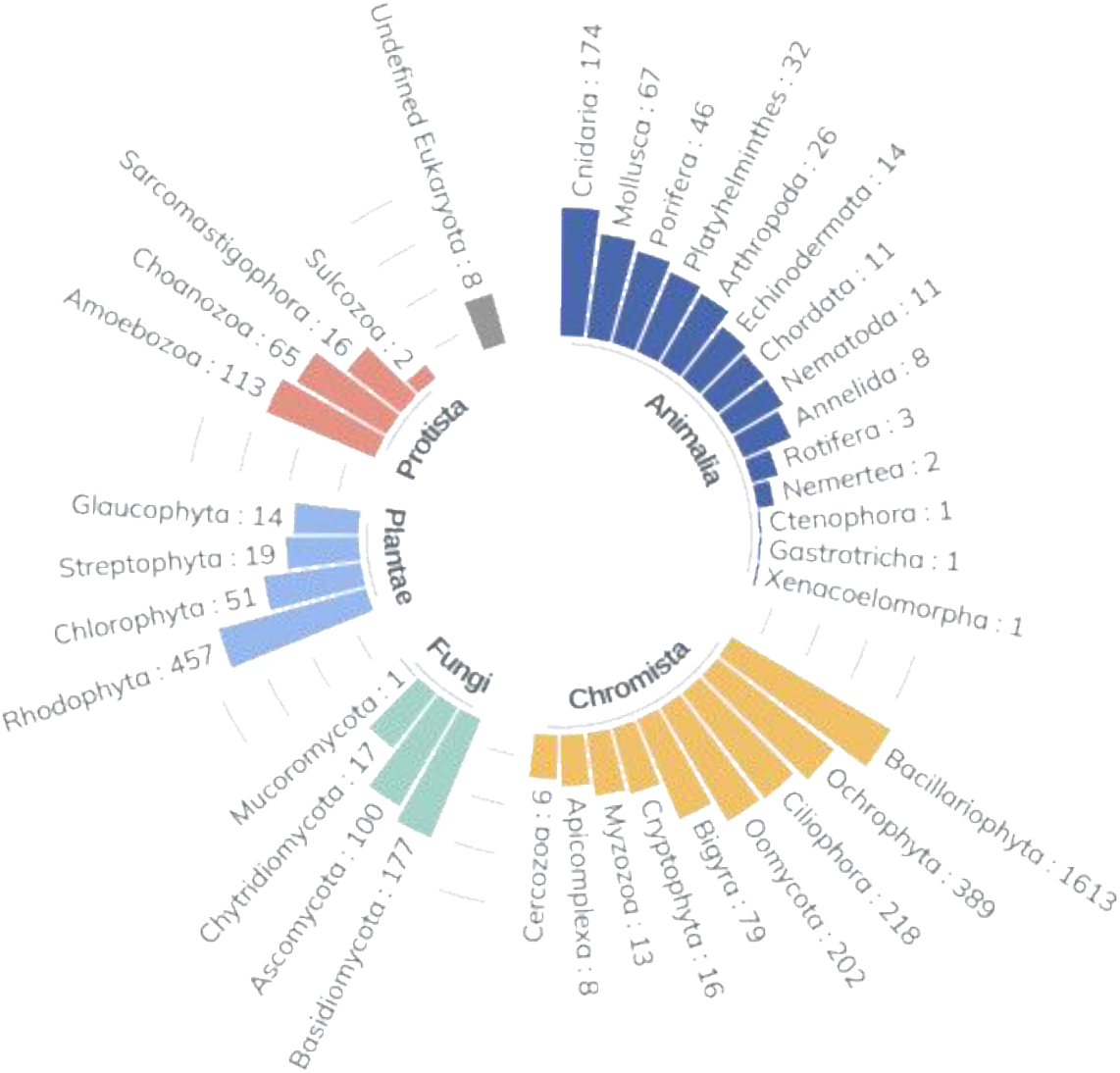
ASVs observed per phyla. Bar size and the number associated with each phylum name indicates the number of unique ASVs assigned to that phylum across all fourteen samples from all three sites. Phyla are grouped by kingdom, following the taxonomic used by the World Register of Marine Species (WoRMS) database. Logarithmic transformation was applied to the scale to enhance visibility.

## Disclaimer/Publisher’s Note

The statements, opinions and data contained in all publications are solely those of the individual author(s) and contributor(s) and not of MDPI and/or the editor(s). MDPI and/or the editor(s) disclaim responsibility for any injury to people or property resulting from any ideas, methods, instructions or products referred to in the content.

## References

1. Gardner, T.A.; Côté, I.M.; Gill, J.A.; Grant, A.; Watkinson, A.R. Long-Term Region-Wide Declines in Caribbean Corals. Science 2003, 301, 958–960, doi:10.1126/science.1086050.

2. Bruno, J.F.; Valdivia, A. Coral Reef Degradation Is Not Correlated with Local Human Population Density. Sci. Rep. 2016, 6, 29778, doi:10.1038/srep29778.

3. Hochberg, E.J.; Gierach, M.M. Missing the Reef for the Corals: Unexpected Trends Between Coral Reef Condition and the Environment at the Ecosystem Scale. Front. Mar. Sci. 2021, 8.

4. Whaley, Z.; Cramer, K.; McClenachan, L.; Tewfik, A.; Alvarez-Filip, L.; McField, M.; Carilli, J.; Vardi, T. Long-Term Change in Caribbean Reef Water Quality and Ecosystem Health. Bull. Fla. Mus. Nat. Hist. 2023, 60, 126–126, doi:10.58782/flmnh.bhpv7556.

5. Grottoli, A.G.; Warner, M.E.; Levas, S.J.; Aschaffenburg, M.D.; Schoepf, V.; McGinley, M.; Baumann, J.; Matsui, Y. The Cumulative Impact of Annual Coral Bleaching Can Turn Some Coral Species Winners into Losers. Glob. Change Biol. 2014, 20, 3823–3833, doi:10.1111/gcb.12658.

6. Kalmus, P.; Ekanayaka, A.; Kang, E.; Baird, M.; Gierach, M. Past the Precipice? Projected Coral Habitability Under Global Heating. Earths Future 2022, 10, e2021EF002608, doi:10.1029/2021EF002608.

7. Reaka, M. The Global Biodiversity of Coral Reefs: A Comparison with Rainforests. 1997.

8. Knowlton, N.; Brainard, R.; Fisher, R.; Moews, M.; Plaisance, L.; Caley, M. Coral Reef Biodiversity. In Life in the World’s Oceans: Diversity, Distribution, and Abundance; 2010; pp. 65–78 ISBN 978-1-4051-9297-2.

9. Battaglia, F.M. Climate Change and the Ocean: The Disruption of the Coral Reef. In; 2023; pp. 121–130 ISBN 978-3-031-24887-0.

10. Edwards, A.; Guest, J.; Humanes, A. Rehabilitating Coral Reefs in the Anthropocene. Curr. Biol. 2024, 34, R399– R406, doi:10.1016/j.cub.2023.12.054.

11. Boström-Einarsson, L.; Babcock, R.C.; Bayraktarov, E.; Ceccarelli, D.; Cook, N.; Ferse, S.C.A.; Hancock, B.; Harrison, P.; Hein, M.; Shaver, E.;, et al. Coral Restoration – A Systematic Review of Current Methods, Successes, Failures and Future Directions. PLOS ONE 2020, 15, e0226631, doi:10.1371/journal.pone.0226631.

12. Higgins, E.; Metaxas, A.; Scheibling, R.E. A Systematic Review of Artificial Reefs as Platforms for Coral Reef Research and Conservation. PLOS ONE 2022, 17, e0261964, doi:10.1371/journal.pone.0261964.

13. Sedano, F.; Navarro-Barranco, C.; Guerra-García, J.M.; Espinosa, F. Understanding the Effects of Coastal Defence Structures on Marine Biota: The Role of Substrate Composition and Roughness in Structuring Sessile, Macro- and Meiofaunal Communities. Mar. Pollut. Bull. 2020, 157, 111334, doi:10.1016/j.marpolbul.2020.111334.

14. Hata, T.; Madin, J.S.; Cumbo, V.R.; Denny, M.; Figueiredo, J.; Harii, S.; Thomas, C.J.; Baird, A.H. Coral Larvae Are Poor Swimmers and Require Fine-Scale Reef Structure to Settle. Sci. Rep. 2017, 7, 2249, doi:10.1038/s41598-017-02402-y.

15. Levy, N.; Berman, O.; Yuval, M.; Loya, Y.; Treibitz, T.; Tarazi, E.; Levy, O. Emerging 3D Technologies for Future Reformation of Coral Reefs: Enhancing Biodiversity Using Biomimetic Structures Based on Designs by Nature. Sci. Total Environ. 2022, 830, 154749, doi:10.1016/j.scitotenv.2022.154749.

16. Spieler, R.E.; Gilliam, D.S.; Sherman, R.L. Artificial Substrate and Coral Reef Restoration: What Do We Need to Know to Know What We Need? Bull. Mar. Sci. 2001, 69, 1013–1030.

17. Vivier, B.; Dauvin, J.-C.; Navon, M.; Rusig, A.-M.; Mussio, I.; Orvain, F.; Boutouil, M.; Claquin, P. Marine Artificial Reefs, a Meta-Analysis of Their Design, Objectives and Effectiveness. Glob. Ecol. Conserv. 2021, 27, e01538, doi:10.1016/j.gecco.2021.e01538.

18. Done, T.J. Phase Shifts in Coral Reef Communities and Their Ecological Significance. In Proceedings of the The Ecology of Mangrove and Related Ecosystems; Jaccarini, V., Martens, E., Eds.; Springer Netherlands: Dordrecht, 1992; pp. 121–132.

19. Wismer, S.; Hoey, A.; Bellwood, D. Cross-Shelf Benthic Community Structure on the Great Barrier Reef: Relationships between Macroalgal Cover and Herbivore Biomass. Mar. Ecol. Prog. Ser. 2009, 376, 45–54, doi:10.3354/meps07790.

20. Edwards, A.; Job, S.; Wells, S. Learning Lessons from Past Reef-Rehabilitation Projects. In Reef rehabilitation manual; 2010; pp. 129–166 ISBN 978-1-921317-05-7.

21. Hein, M.Y.; Willis, B.L.; Beeden, R.; Birtles, A. The Need for Broader Ecological and Socioeconomic Tools to Evaluate the Effectiveness of Coral Restoration Programs. Restor. Ecol. 2017, 25, 873–883, doi:10.1111/rec.12580.

22. Bourne, D.G.; Morrow, K.M.; Webster, N.S. Insights into the Coral Microbiome: Underpinning the Health and Resilience of Reef Ecosystems. Annu. Rev. Microbiol. 2016, 70, 317–340, doi:10.1146/annurev-micro-102215-095440.

23. Leray, M.; Knowlton, N. DNA Barcoding and Metabarcoding of Standardized Samples Reveal Patterns of Marine Benthic Diversity. Proc. Natl. Acad. Sci. 2015, 112, 2076–2081, doi:10.1073/pnas.1424997112.

24. Beng, K.C.; Corlett, R.T. Applications of Environmental DNA (eDNA) in Ecology and Conservation: Opportunities, Challenges and Prospects. Biodivers. Conserv. 2020, 29, 2089–2121, doi:10.1007/s10531-020-01980-0.

25. Thompson, S.; Jarman, S.; Griffin, K.; Spencer, C.; Cummins, G.; Partridge, J.; Langlois, T. Novel Drop-Sampler for Simultaneous Collection of Stereo-Video, Environmental DNA and Oceanographic Data. Ecol. Evol. 2024, 14, doi:10.1002/ece3.70705.

26. Duprey, N.N.; McIlroy, S.E.; Ng, T.P.T.; Thompson, P.D.; Kim, T.; Wong, J.C.Y.; Wong, C.W.M.; Husa, S.M.; Li, S.M.H.; Williams, G.A.;, et al. Facing a Wicked Problem with Optimism: Issues and Priorities for Coral Conservation in Hong Kong. Biodivers. Conserv. 2017, 26, 2521–2545, doi:10.1007/s10531-017-1383-z.

27. Xie, J.Y.; Yeung, Y.H.; Kwok, C.K.; Kei, K.; Ang, P.; Chan, L.L.; Cheang, C.C.; Chow, W.; Qiu, J.-W. Localized Bleaching and Quick Recovery in Hong Kong’s Coral Communities. Mar. Pollut. Bull. 2020, 153, 110950, doi:10.1016/j.marpolbul.2020.110950.

28. Duprey, N.N.; Yasuhara, M.; Baker, D.M. Reefs of Tomorrow: Eutrophication Reduces Coral Biodiversity in an Urbanized Seascape. Glob. Change Biol. 2016, 22, 3550–3565.

29. Fabricius, K.E. Effects of Terrestrial Runoff on the Ecology of Corals and Coral Reefs: Review and Synthesis. Mar. Pollut. Bull. 2005, 50, 125–146, doi:10.1016/j.marpolbul.2004.11.028.

30. Cybulski, J.D.; Husa, S.M.; Duprey, N.N.; Mamo, B.L.; Tsang, T.P.; Yasuhara, M.; Xie, J.Y.; Qiu, J.-W.; Yokoyama, Y.; Baker, D.M. Coral Reef Diversity Losses in China’s Greater Bay Area Were Driven by Regional Stressors. Sci. Adv. 2020, 6, eabb1046.

31. Yeung, Y.H.; Xie, J.Y.; Kwok, C.K.; Kei, K.; Ang, P.; Chan, L.L.; Dellisanti, W.; Cheang, C.C.; Chow, W.K.; Qiu, J.-W. Hong Kong’s Subtropical Scleractinian Coral Communities: Baseline, Environmental Drivers and Management Implications. Mar. Pollut. Bull. 2021, 167, 112289, doi:10.1016/j.marpolbul.2021.112289.

32. Hua, F.L.; Tsang, Y.F.; Chua, H. Progress of Water Pollution Control in Hong Kong. Aquat. Ecosyst. Health Manag. 2008, 11, 225–229, doi:10.1080/14634980802100717.

33. Lange, C. Rethinking Artificial Reef Structures through a Robotic 3D Clay Printing Method. In Proceedings of the Proceedings of the 25th International Conference of the Association for Computer-Aided Architectural Design Research in Asia (CAADRIA); Association for Computer-Aided Architectural Design Research in Asia (CAADRIA): Hong Kong, 2020; Vol. 2, pp. 463–472.

34. Brooks, M. E.; Kristensen, K.; Benthem, K. J. van; Magnusson, A.; Berg, C. W.; Nielsen, A.; Skaug, H. J.; Mächler, M.; Bolker, B. M. glmmTMB Balances Speed and Flexibility Among Packages for Zero-Inflated Generalized Linear Mixed Modeling. R J. 2017, 9, 378, doi:10.32614/RJ-2017-066.

35. Bates, D.; Mächler, M.; Bolker, B.; Walker, S. Fitting Linear Mixed-Effects Models Using Lme4. J. Stat. Softw. 2015, 67, 1–48, doi:10.18637/jss.v067.i01.

36. Lenth, R. Estimated Marginal Means, Aka Least-Squares Means 2024.

37. Hodgson, G. Reef Check California Instruction Manual: A Guide to Monitoring California’s Rocky Reefs; 1st ed.; Reef Check Foundation: Pacific Palisades, CA, 2006; ISBN 978-0-9723051-9-8.

38. Bolyen, E.; Rideout, J.R.; Dillon, M.R.; Bokulich, N.A.; Abnet, C.C.; Al-Ghalith, G.A.; Alexander, H.; Alm, E.J.; Arumugam, M.; Asnicar, F.;, et al. Reproducible, Interactive, Scalable and Extensible Microbiome Data Science Using QIIME 2. Nat. Biotechnol. 2019, 37, 852–857, doi:10.1038/s41587-019-0209-9.

39. Callahan, B.J.; McMurdie, P.J.; Rosen, M.J.; Han, A.W.; Johnson, A.J.A.; Holmes, S.P. DADA2: High-Resolution Sample Inference from Illumina Amplicon Data. Nat. Methods 2016, 13, 581–583, doi:10.1038/nmeth.3869.

40. Wang, S.; Meyer, E.; McKay, J.K.; Matz, M.V. 2b-RAD: A Simple and Flexible Method for Genome-Wide Genotyping. Nat. Methods 2012, 9, 808–810, doi:10.1038/nmeth.2023.

41. Porter, T.M.; Hajibabaei, M. Over 2.5 Million COI Sequences in GenBank and Growing. PLOS ONE 2018, 13, e0200177, doi:10.1371/journal.pone.0200177.

42. Team, R.C.; Team, M.R.C.; Suggests, M.; Matrix, S. Package Stats. R Stats Package 2018, 1–3.

43. McMurdie, P.J.; Holmes, S. Phyloseq: An R Package for Reproducible Interactive Analysis and Graphics of Microbiome Census Data. PloS One 2013, 8, e61217.

44. Kandlikar, G.S.; Gold, Z.J.; Cowen, M.C.; Meyer, R.S.; Freise, A.C.; Kraft, N.J.; Moberg-Parker, J.; Sprague, J.; Kushner, D.J.; Curd, E.E. Ranacapa: An R Package and Shiny Web App to Explore Environmental DNA Data with Exploratory Statistics and Interactive Visualizations. F1000Research 2018, 7, 1734.

45. Wickham, H. Getting Started with Ggplot2. In ggplot2: Elegant graphics for data analysis; Springer, 2016; pp. 11–31.

46. Oksanen, J.; Blanchet, F.G.; Kindt, R.; Legendre, P.; Minchin, P.R.; O’hara, R.; Simpson, G.L.; Solymos, P.; Stevens, M.H.H.; Wagner, H. Package ‘Vegan.’ Community Ecol. Package Version 2013, 2, 1–295.

47. Kassambara, A. Rstatix: Pipe-Friendly Framework for Basic Statistical Tests. CRAN Contrib. Packag. 2019.

48. Martinez Arbizu, P. pairwiseAdonis: Pairwise Multilevel Comparison Using Adonis. R Package Version 04 2020, 1.

49. Goergen, E.A.; Schopmeyer, S.; Moulding, A.L.; Moura, A.; Kramer, P.; Viehman, T.S. Coral Reef Restoration Monitoring Guide: Methods to Evaluate Restoration Success from Local to Ecosystem Scales. 2020, doi:10.25923/xndz-h538.

50. Gladfelter, E.H.; Monahan, R.K.; Gladfelter, W.B. Growth Rates of Five Reef-Building Corals in the Northeastern Caribbean. Bull. Mar. Sci. 1978, 28, 728–734.

51. Zawada, K.J.; Dornelas, M.; Madin, J.S. Quantifying Coral Morphology. Coral Reefs 2019, 38, 1281–1292.

52. Rossi, S.; Schubert, N.; Brown, D.; Soares, M. de O.; Grosso, V.; Rangel-Huerta, E.; Maldonado, E. Linking Host Morphology and Symbiont Performance in Octocorals. Sci. Rep. 2018, 8, 12823.

53. Lough, J.; Barnes, D. Environmental Controls on Growth of the Massive Coral Porites. J. Exp. Mar. Biol. Ecol. 2000, 245, 225–243.

54. Browne, N. Spatial and Temporal Variations in Coral Growth on an Inshore Turbid Reef Subjected to Multiple Disturbances. Mar. Environ. Res. 2012, 77, 71–83.

55. Calle-Triviño, J.; Muñiz-Castillo, A.I.; Cortés-Useche, C.; Morikawa, M.; Sellares-Blasco, R.; Arias-González, J.E. Approach to the Functional Importance of Acropora Cervicornis in Outplanting Sites in the Dominican Republic. Front. Mar. Sci. 2021, 8, 668325.

56. Young, C.N.; Schopmeyer, S.; Lirman, D. A Review of Reef Restoration and Coral Propagation Using the Threatened Genus Acropora in the Caribbean and Western Atlantic. Bull. Mar. Sci. 2012, 88, 1075–1098, doi:10.5343/BMS.2011.1143.

57. Highsmith, R.C. Reproduction by Fragmentation in Corals. Mar. Ecol. Prog. Ser. Oldendorf 1982, 7, 207–226.

58. Lirman, D. Fragmentation in the Branching Coral Acropora Palmata (Lamarck): Growth, Survivorship, and Reproduction of Colonies and Fragments. J. Exp. Mar. Biol. Ecol. 2000, 251, 41–57, doi:10.1016/S0022-0981(00)00205-7.

59. Clark, S.; Edwards, A. Coral Transplantation as an Aid to Reef Rehabilitation: Evaluation of a Case Study in the Maldive Islands. Coral Reefs 1995, 14, 201–213.

60. Yap, H.T.; Alino, P.M.; Gomez, E.D. Trends in Growth and Mortality of Three Coral Species(Anthozoa: Scleractinia), Including Effects of Transplantation. Mar. Ecol. Prog. Ser. Oldendorf 1992, 83, 91–101.

61. Ware, M.; Garfield, E.N.; Nedimyer, K.; Levy, J.; Kaufman, L.; Precht, W.; Winters, R.S.; Miller, S.L. Survivorship and Growth in Staghorn Coral (Acropora Cervicornis) Outplanting Projects in the Florida Keys National Marine Sanctuary. PLoS One 2020, 15, e0231817.

62. Cabaitan, P.C.; Yap, H.T.; Gomez, E.D. Performance of Single versus Mixed Coral Species for Transplantation to Restore Degraded Reefs. Restor. Ecol. 2015, 23, 349–356, doi:10.1111/rec.12205.

63. Conti-Jerpe, I.E.; Thompson, P.D.; Wong, C.W.M.; Oliveira, N.L.; Duprey, N.N.; Moynihan, M.A.; Baker, D.M. Trophic Strategy and Bleaching Resistance in Reef-Building Corals. Sci. Adv. 2020, 6, eaaz5443, doi:10.1126/sciadv.aaz5443.

64. Cybulski, J.D. Hong Kong’s Coral Assemblages through Time : A Paleoecological and Geochemical Look at Human-Driven Change, The University of Hong Kong: Pokfulam, Hong Kong SAR., 2021.

65. Bohnsack, J.A.; Sutherland, D.L. Artificial Reef Research: A Review with Recommendations for Future Priorities. Bull. Mar. Sci. 1985, 37, 11–39.

66. Arena, P.T.; Jordan, L.K.B.; Spieler, R.E. Fish Assemblages on Sunken Vessels and Natural Reefs in Southeast Florida, USA. In Proceedings of the Biodiversity in Enclosed Seas and Artificial Marine Habitats; Relini, G., Ryland, J., Eds.; Springer Netherlands: Dordrecht, 2007; pp. 157–171.

67. Gratwicke, B.; Speight, M.R. The Relationship between Fish Species Richness, Abundance and Habitat Complexity in a Range of Shallow Tropical Marine Habitats. J. Fish Biol. 2005, 66, 650–667, doi:10.1111/j.0022-1112.2005.00629.x.

68. Santos, L.N.; Araujo, F.G.; Brotto, D.S. Artificial Structures as Tools for Fish Habitat Rehabilitation in a Neotropical Reservoir. Aquat. Conserv. Mar. Freshw. Ecosyst. 2008, 18, 896.

69. Sherman, R.L.; Gilliam, D.S.; Spieler, R.E. Artificial Reef Design: Void Space, Complexity, and Attractants. ICES J. Mar. Sci. 2002, 59, S196–S200.

70. Burkepile, D.E.; Hay, M.E. Herbivore Species Richness and Feeding Complementarity Affect Community Structure and Function on a Coral Reef. Proc. Natl. Acad. Sci. 2008, 105, 16201–16206, doi:10.1073/pnas.0801946105.

71. Hughes, T.P.; Rodrigues, M.J.; Bellwood, D.R.; Ceccarelli, D.; Hoegh-Guldberg, O.; McCook, L.; Moltschaniwskyj, N.; Pratchett, M.S.; Steneck, R.S.; Willis, B. Phase Shifts, Herbivory, and the Resilience of Coral Reefs to Climate Change. Curr. Biol. 2007, 17, 360–365, doi:10.1016/j.cub.2006.12.049.

72. Mumby, P.; Steneck, R. Coral Reef Management and Conservation in Light of Rapidly Evolving Ecological Paradigms. Trends Ecol. Evol. 2008, 23, 555–563, doi:10.1016/j.tree.2008.06.011.

73. Glynn, P.W.; D’Croz, L. Experimental Evidence for High Temperature Stress as the Cause of El Niño-Coincident Coral Mortality. Coral Reefs 1990, 8, 181–191, doi:10.1007/BF00265009.

74. Carreiro-Silva, M.; McClanahan, T.R. Macrobioerosion of Dead Branching Porites, 4 and 6 Years after Coral Mass Mortality. Mar. Ecol. Prog. Ser. 2012, 458, 103–122.

75. Van Der Plas, F. Biodiversity and Ecosystem Functioning in Naturally Assembled Communities. Biol. Rev. 2019, 94, 1220–1245, doi:10.1111/brv.12499.

76. MacNeil, M.A.; Graham, N.A.J.; Conroy, M.J.; Fonnesbeck, C.J.; Polunin, N.V.C.; Rushton, S.P.; Chabanet, P.; McClanahan, T.R. Detection Heterogeneity in Underwater Visual-census Data. J. Fish Biol. 2008, 73, 1748–1763, doi:10.1111/j.1095-8649.2008.02067.x.

77. Yachi, S.; Loreau, M. Biodiversity and Ecosystem Productivity in a Fluctuating Environment: The Insurance Hypothesis. Proc. Natl. Acad. Sci. 1999, 96, 1463–1468, doi:10.1073/pnas.96.4.1463.

78. McGrady-Steed, J.; Harris, P.M.; Morin, P.J. Biodiversity Regulates Ecosystem Predictability. Nature 1997, 390, 162– 165.

79. Naeem, S.; Li, S. Biodiversity Enhances Ecosystem Reliability. Nature 1997, 390, 507–509, doi:10.1038/37348.

80. Gann, G.D.; McDonald, T.; Walder, B.; Aronson, J.; Nelson, C.R.; Jonson, J.; Hallett, J.G.; Eisenberg, C.; Guariguata, M.R.; Liu, J. International Principles and Standards for the Practice of Ecological Restoration. Restor. Ecol. 2019, 27, S1–S46.

81. McDonald, T.; Gann, G.; Jonson, J.; Dixon, K. International Standards for the Practice of Ecological Restoration– Including Principles and Key Concepts.(Society for Ecological Restoration: Washington, DC, USA.). Soil-Tec Inc© Marcel Huijser Bethanie Walder 2016.

82. Perkol-Finkel, S.; Benayahu, Y. Recruitment of Benthic Organisms onto a Planned Artificial Reef: Shifts in Community Structure One Decade Post-Deployment. Mar. Environ. Res. 2005, 59, 79–99, doi:10.1016/j.marenvres.2004.03.122.

83. Spagnolo, A.; Cuicchi, C.; Punzo, E.; Santelli, A.; Scarcella, G.; Fabi, G. Patterns of Colonization and Succession of Benthic Assemblages in Two Artificial Substrates. J. Sea Res. 2014, 88, 78–86, doi:10.1016/j.seares.2014.01.007.

84. Pickett, S.T.A.; Cadenasso, M.L.; Rosi-Marshall, E.J.; Belt, K.T.; Groffman, P.M.; Grove, J.M.; Irwin, E.G.; Kaushal, S.S.; LaDeau, S.L.; Nilon, C.H.;, et al. Dynamic Heterogeneity: A Framework to Promote Ecological Integration and Hypothesis Generation in Urban Systems. Urban Ecosyst. 2017, 20, 1–14, doi:10.1007/s11252-016-0574-9.

85. Todd, P.A.; Heery, E.C.; Loke, L.H.L.; Thurstan, R.H.; Kotze, D.J.; Swan, C. Towards an Urban Marine Ecology: Characterizing the Drivers, Patterns and Processes of Marine Ecosystems in Coastal Cities. Oikos 2019, 128, 1215– 1242, doi:10.1111/oik.05946.

86. Bae, S.; Ubagan, M.D.; Shin, S.; Kim, D.G. Comparison of Recruitment Patterns of Sessile Marine Invertebrates According to Substrate Characteristics. Int. J. Environ. Res. Public. Health 2022, 19, 1083, doi:10.3390/ijerph19031083.

87. Juan, S. de; Thrush, S.F.; Hewitt, J.E. Counting on β-Diversity to Safeguard the Resilience of Estuaries. PLOS ONE 2013, 8, e65575, doi:10.1371/journal.pone.0065575.

88. Oliver, T.H.; Heard, M.S.; Isaac, N.J.B.; Roy, D.B.; Procter, D.; Eigenbrod, F.; Freckleton, R.; Hector, A.; Orme, C.D.L.; Petchey, O.L.;, et al. Biodiversity and Resilience of Ecosystem Functions. Trends Ecol. Evol. 2015, 30, 673–684, doi:10.1016/j.tree.2015.08.009.

89. Connell, S.D. Floating Pontoons Create Novel Habitats for Subtidal Epibiota. J. Exp. Mar. Biol. Ecol. 2000, 247, 183– 194, doi:10.1016/S0022-0981(00)00147-7.

90. Perkol-Finkel, S.; Benayahu, Y. Differential Recruitment of Benthic Communities on Neighboring Artificial and Natural Reefs. J. Exp. Mar. Biol. Ecol. 2007, 340, 25–39, doi:10.1016/j.jembe.2006.08.008.

91. McIlroy, S.E.; Guibert, I.; Archana, A.; Chung, W.Y.H.; Duffy, J.E.; Gotama, R.; Hui, J.; Knowlton, N.; Leray, M.; Meyer, C.;, et al. Life Goes on: Spatial Heterogeneity Promotes Biodiversity in an Urbanized Coastal Marine Ecosystem. Glob. Change Biol. 2023, 30, e17248, doi:10.1111/gcb.17248.

92. Atkinson, J.; Brudvig, L.A.; Mallen-Cooper, M.; Nakagawa, S.; Moles, A.T.; Bonser, S.P. Terrestrial Ecosystem Restoration Increases Biodiversity and Reduces Its Variability, but Not to Reference Levels: A Global Meta-analysis. Ecol. Lett. 2022, 25, 1725–1737, doi:10.1111/ele.14025.

93. Bayraktarov, E.; Stewart-Sinclair, P.J.; Brisbane, S.; Boström-Einarsson, L.; Saunders, M.I.; Lovelock, C.E.; Possingham, H.P.; Mumby, P.J.; Wilson, K.A. Motivations, Success, and Cost of Coral Reef Restoration. Restor. Ecol. 2019, 27, 981–991, doi:10.1111/rec.12977.

94. Sato, M.; Inoue, N.; Nambu, R.; Furuichi, N.; Imaizumi, T.; Ushio, M. Quantitative Assessment of Multiple Fish Species around Artificial Reefs Combining Environmental DNA Metabarcoding and Acoustic Survey. Sci. Rep. 2021, 11, 19477.

95. Inoue, N.; Sato, M.; Furuichi, N.; Imaizumi, T.; Ushio, M. The Relationship between eDNA Density Distribution and Current Fields around an Artificial Reef in the Waters of Tateyama Bay, Japan. Metabarcoding Metagenomics 2022, 6, e87415.

96. Krolow, A.P. Assessing the Diversity of Fish Communities at or Around Artificial Reefs Along the Louisiana Coast Through the Use of Environmental DNA (eDNA). 2019.

97. Krolow, A.D.; Geheber, A.D.; Piller, K.R. If You Build It, Will They Come? An Environmental DNA Assessment of Fish Assemblages on Artificial Reefs in the Northern Gulf of Mexico. Trans. Am. Fish. Soc. 2022, 151, 297–321.

98. Miyajima-Taga, Y.; Sato, M.; Oi, K.; Furuichi, N.; Inoue, N. Fine-Scale Spatial Distribution of a Fish Community in Artificial Reefs Investigated Using an Underwater Drone and Environmental DNA Analysis. Mar. Ecol. Prog. Ser. 2024, 740, 123–144.

99. Levy, N.; Simon-Blecher, N.; Ben-Ezra, S.; Yuval, M.; Doniger, T.; Leray, M.; Karako-Lampert, S.; Tarazi, E.; Levy, O. Evaluating Biodiversity for Coral Reef Reformation and Monitoring on Complex 3D Structures Using Environmental DNA (eDNA) Metabarcoding. Sci. Total Environ. 2023, 856, 159051, doi:10.1016/j.scitotenv.2022.159051.

100. Knoester, E.; Rienstra, J.; Schürmann, Q.; Wolma, A.; Murk, A.; Osinga, R. Community-Managed Coral Reef Restoration in Southern Kenya Initiates Reef Recovery Using Various Artificial Reef Designs. Front. Mar. Sci. 2023, 10, 1152106.

101. Harrington, L.; Fabricius, K.; De’ath, G.; Negri, A. Recognition and Selection of Settlement Substrata Determine Post-Settlement Survival in Corals. Ecology 2004, 85, 3428–3437, doi:10.1890/04-0298.

102. Webster, N.S.; Soo, R.; Cobb, R.; Negri, A.P. Elevated Seawater Temperature Causes a Microbial Shift on Crustose Coralline Algae with Implications for the Recruitment of Coral Larvae. ISME J. 2011, 5, 759–770, doi:10.1038/ismej.2010.152.

103. Bessey, C.; Jarman, S.N.; Berry, O.; Olsen, Y.S.; Bunce, M.; Simpson, T.; Power, M.; McLaughlin, J.; Edgar, G.J.; Keesing, J. Maximizing Fish Detection with eDNA Metabarcoding. *Environ*. DNA 2020, 2, 493–504.

